# Automatic segmentation of *Drosophila* neural compartments using GAL4 expression data reveals novel visual pathways

**DOI:** 10.1101/032292

**Authors:** K Panser, L Tirian, F Schulze, S Villalba, GSXE Jefferis, Katja Bühler, AD Straw

## Abstract

We made use of two recent, large-scale *Drosophila* GAL4 libraries and associated confocal imaging datasets to automatically segment large brain regions into smaller putative functional units such as neuropils and fiber tracts. The method we developed is based on the hypothesis that molecular identity can be used to assign individual voxels to biologically meaningful regions. Our results (available at https://strawlab.org/braincode) are consistent with this hypothesis because regions with well-known anatomy, namely the antennal lobes and central complex, were automatically segmented into familiar compartments. We then applied the algorithm to the central brain regions receiving input from the optic lobes. Based on the automated segmentation and manual validation, we can identify and provide promising driver lines for 10 previously identified and 14 novel types of visual projection neurons and their associated optic glomeruli. The same strategy can be used in other brain regions and likely other species, including vertebrates.

## Introduction

A key goal of neuroscientists is to understand brain function through a mechanistic understanding of the physiology and anatomy of circuits within the brain and their relation to behavior. Recently developed neurogenetic tools allowing genetic targeting of specific cell classes and brain regions have been essential to many advances in the past couple decades. More recently, large-scale efforts to develop collections of thousands of *Drosophila* lines in which GAL4 expression is controlled via fragments of genomic DNA containing putative enhancers and repressors (Jenett et al., 2012; Kvon et al., 2014; Pfeiffer et al., 2008) have already been productively used as the basis for numerous screens, targeted neuronal manipulation, and anatomical studies.

For many regions of the brain, we lack both a detailed anatomical understanding of the structures present and the ability to reproducibly target specific cell types contained within those structures with genetic tools. For example, despite extensive work on the visual system of flies such as *Drosophila* (Fischbach and Dittrich, 1989; Fischbach and Lyly-Hünerberg, 1983; Nern et al., 2015; Raghu et al., 2011, 2009, 2007; Raghu and Borst, 2011), the major targets of visual projection neurons (VPNs), cells whose projections leave the optic lobes and target regions of the central brain, remain relatively uncharacterized despite several pioneering papers (Aptekar et al., 2015; Fischbach and Dittrich, 1989; Fischbach and Lyly-Hünerberg, 1983; Ito et al., 2013; Mu et al., 2012; Okamura and Strausfeld, 2007; Otsuna et al., 2014; Otsuna and Ito, 2006; Strausfeld et al., 2007; Strausfeld and Bacon, 1983; Strausfeld and Lee, 1991; Strausfeld and Okamura, 2007). This region is particularly interesting because the VPNs are an information bottleneck; visual information must pass through the VPNs before it can influence behavior and the numbers of cell types and cell numbers are small. For example, in the stalk-eyed fly *Cytrodiopsis whitei*, the optic nerve contains about 6000 axons (Burkhardt and Motte, 1983) and the number of VPN types in *Drosophila* is thought to number about 50 (Otsuna and Ito, 2006). Typically, many of a single VPN type will converge onto a glomerular structure (Strausfeld and Bacon, 1983; Strausfeld and Lee, 1991). The suggestion is that these optic glomeruli may process visual features in a way analogous to olfactory glomeruli in the antennal lobe (Mu et al., 2012) although the visual projection neurons are likely four or five synapses from the neurons involved in sensory transduction while the olfactory glomeruli are the primary processing centers to which the olfactory sensory neurons converge. As it has been with the *Drosophila* olfactory system, genetic access to the VPN cell types and other cell types innervating the optic glomeruli will be useful in elucidating visual circuit function.

Similarly, other regions of ‘terra incognita,’ brain regions which remain largely undescribed, exist both within fly and vertebrate, including human, brains (Alkemade et al., 2013; Ito et al., 2013), and an automatic approach to discover functional units, such as nuclei or axon tracts, and to suggest candidate genetic lines that could be used for specific targeting of these regions would be useful. Indeed – apart from the antennal lobes, mushroom bodies, and central complex – much of the *Drosophila* brain appears homogeneous with conventional histological techniques (Ito et al., 2013). Several projects have made use of clonal analyses in which rare stochastic genetic events isolate a small number of neurons and consequently assembling many such examples allows detailed reconstructions of specific cell types and hypotheses about brain structures (Chiang et al., 2011; Hadjieconomou et al., 2011; Hampel et al., 2011; Ito et al., 2013; Livet et al., 2007; Shih et al., 2015; Yu et al., 2013). Other efforts combine electron microscopy with serial reconstruction to produce even more detailed connectomic data (Cardona et al., 2010; Helmstaedter et al., 2013; Takemura et al., 2013; White et al., 1986). Despite their utility at revealing brain structure, these approaches rely on stochastic events or histological techniques that are difficult to correlate with cell-type specific genetically encoded markers and thus the results cannot be directly used to identify promising driver lines for subsequent study.

In this study, we used imaging data from recent *Drosophila* GAL4 collections to automatically identify structure within the fly brain and to identify driver lines targeting these regions. Our approach was based on the hypothesis that multiple locations within a particular nucleus, glomerulus, or axon tract would have patterns of genetic activity, such as gene expression or enhancer activation, more similar to each other than to locations within other structures. RNA expression patterns in mouse (Fakhry and Ji, 2015; Lein et al., 2007; Ng et al., 2009; Thompson et al., 2014) and human brains (Goel et al., 2014; Hawrylycz et al., 2012; Mahfouz et al., 2015; Myers et al., 2015) show this to be true at a relatively course spatial scale – sets of genes expressed in, for example, cortex or cerebellum, are characteristic for those regions across different individuals. Given that enhancers have more specific expression patterns than the genes that they regulate (Kvon et al., 2014), we hypothesized that use of enhancers, rather than genes, would enable parcellation of brain regions on a smaller scale. By clustering GFP signal driven by enhancer-containing genomic fragments, we identified putative functional units. Our results show that, indeed, patterns of genomic-fragment driven expression can be used to automatically extract brain structure. We found that much of the known structure of the well-understood *Drosophila* antennal lobes is automatically found by our method. We further show that this method predicts multiple optic glomeruli and that extensive manual validation with more classical techniques confirms the existence and shape of these structural elements. By using GAL4 collections rather than either spatial profiling of expression patterns from *in situ* hybridization, stochastic genetic strategies or electron microscopic based reconstruction, this approach highlights existing genetic driver lines likely to be useful for studies of localized neural function.

## Results

### Segmentation based on patterns of genomic fragment coexpression

Our approach to segment brain regions into putative ‘functional units’ (nuclei or glomeruli and axon tracts) is based on the idea that multiple locations within such a structure – a brain nucleus, glomerulus, or axon tract, for example – are closer to each other in terms of molecular identity than locations within other structures. We made use of the large imaging datasets from recent *Drosophila* genomic fragment GAL4 collections, and the overall strategy was to use a conventional clustering technique on GAL4-driven expression data to parcellate a brain region (e.g. antennal lobe or lateral protocerebrum) into a number of smaller putative functional units (e.g. individual olfactory or optic glomeruli) based on their genetic code. Because the strategy links the nucleotide sequence within genomic fragments to specific brain regions, we named it ‘Braincode’ and the results can be interactively viewed at https://strawlab.org/braincode.

As input, we took confocal image stacks from the Rubin lab Janelia FlyLight collection (Jenett et al., 2012; Pfeiffer et al., 2008) and from the Dickson lab Vienna Tiles collection (B. Dickson, personal communication). In total, we used data from 3462 Janelia FlyLight and 6022 Vienna Tiles GAL4 driver lines crossed with *UAS-mCD8::GFP*. Each dataset came registered to a dataset-specific template brain with registration error estimated to be 2-3 μm (Cachero et al., 2010; Yu et al., 2010). On a per-voxel basis we calculated the set of driver lines for which GFP expression was higher than a threshold. We used the Dice coefficient to quantify expression similarity between each possible pair of voxels and this *n* x *n* distance matrix was used to group voxels into clusters of similar expression using *k*-medoids clustering (Figure 1, see Methods for details). As typical for clustering algorithms, one parameter controls the number of clusters, and in our case we chose several different values for *k* and evaluated results for different choices and in each of the two independent datasets. Neither manual inspection nor calculation of a metric designed to measure clustering repeatability, adjusted Rand index (Figure 1–figure supplement 1), showed an obvious optimal value for *k*. Therefore, we chose a value of *k* equal 60 as a number which appeared to provide sufficiently many clusters to capture important structures at a small scale without producing an overwhelming number. The result of the clustering algorithm is the assignment of each voxel in the input brain region to one of the *k* clusters. This approach therefore divides the brain into distinct regions, each likely innervated by multiple cell types. While local interneurons might be confined specifically to the region of a particular cluster, other cell types may extend through multiple clusters and into more distant brain regions. The clusters found in this way are predictions of functional units in the *Drosophila* brain. Most of our subsequent efforts were to evaluate the quality of these results.

**Figure 1.**
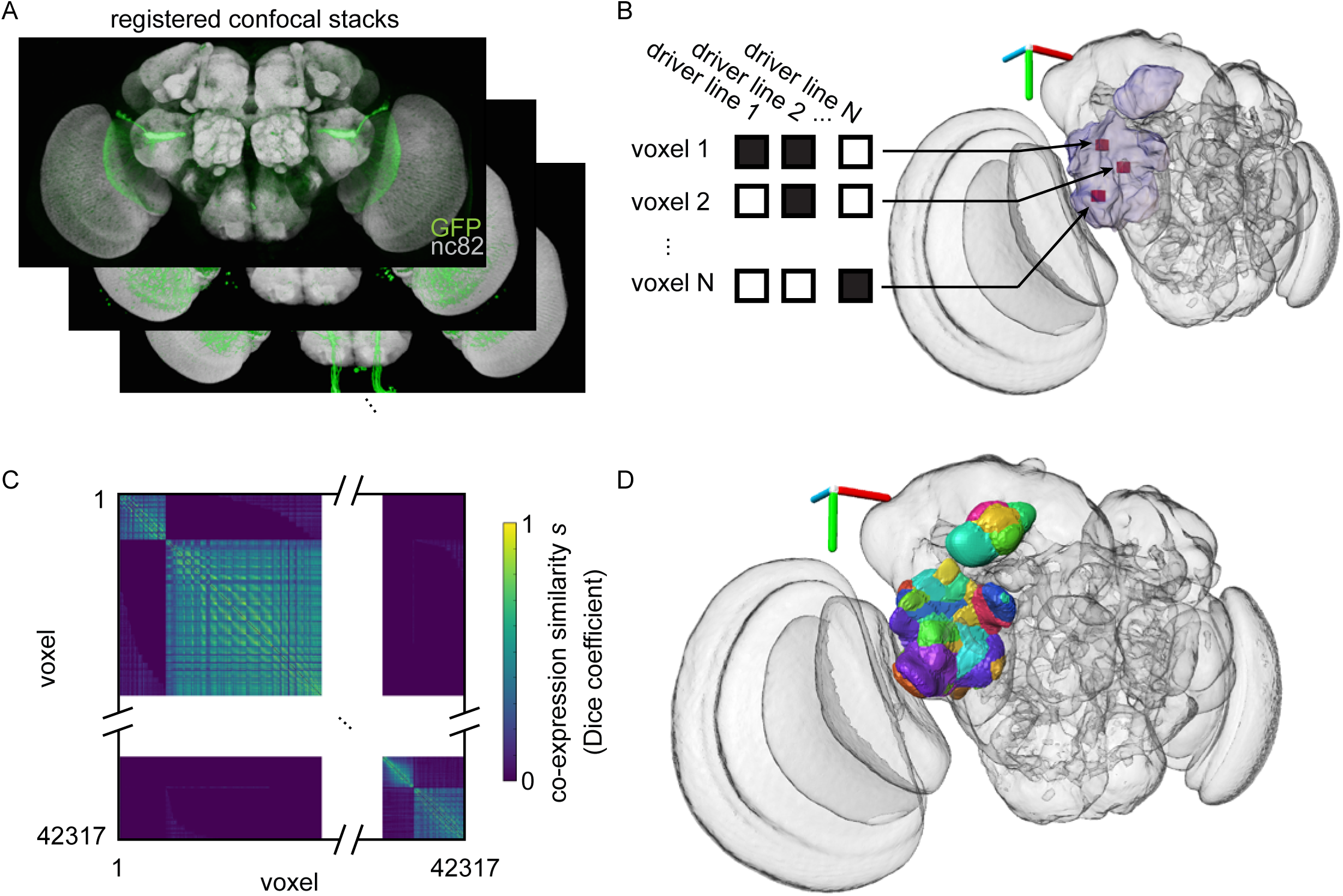
Automatic segmentation of a brain region into domains sharing common enhancer profiles. A) Thousands of registered confocal image stacks from the Janelia FlyLight and Vienna Tiles projects were used. B) Within an analyzed brain region (purple outline), a list of driver lines driving expression was compiled for each voxel. C) A voxel-to-voxel similarity *s* was computed using the Dice coefficient and *k*-medoids was used to cluster groups of voxels of putative functional units. D) Each voxel is colored according to its cluster and plotted in the original brain coordinate system. All panels: Janelia FlyLight data for the optic Ventrolateral Neuropil (oVLNP) region defined as PLP, PVLP, and AOTU, run 1, 42317 voxels, 3462 driver lines, *k* equal 60. 3D axes scale 40 μm in lateral (red), dorsal-ventral (green), anterior-posterior (blue).

If our hypothesis is correct that functional units can be automatically segmented using patterns of coexpression, we can make several predictions. First, despite physical distance not being used as a parameter in defining the clusters, we would expect valid clusters to be spatially compact rather than consisting of, for example, individual voxels scattered throughout the volume. Second, we would expect that for a bilaterally symmetric brain, a given cluster should consist of voxels in mirror-symmetric positions. Third, when clustering is used to segment regions that are already well-understood, the shape, size and location of the automatically found clusters match the known structures. Fourth, when clustering is performed on a different dataset (e.g. Janelia FlyLight versus Vienna Tiles), we expect similar segmentations because the underlying molecular identity of the functional units should dominate the results.

### Automatic segmentation of the antennal lobes

To test these expectations, we examined the Braincode results from the antennal lobe (AL) and central complex (CX) (Figure 2). As shown when run with the number of clusters *k* set to 60, the resulting clusters were compact shapes similar in appearance to the known olfactory glomeruli (Couto et al., 2005; Grabe et al., 2015; Vosshall et al., 2000) filling the volume of the AL (Figure 2A-B).

**Figure 2.**
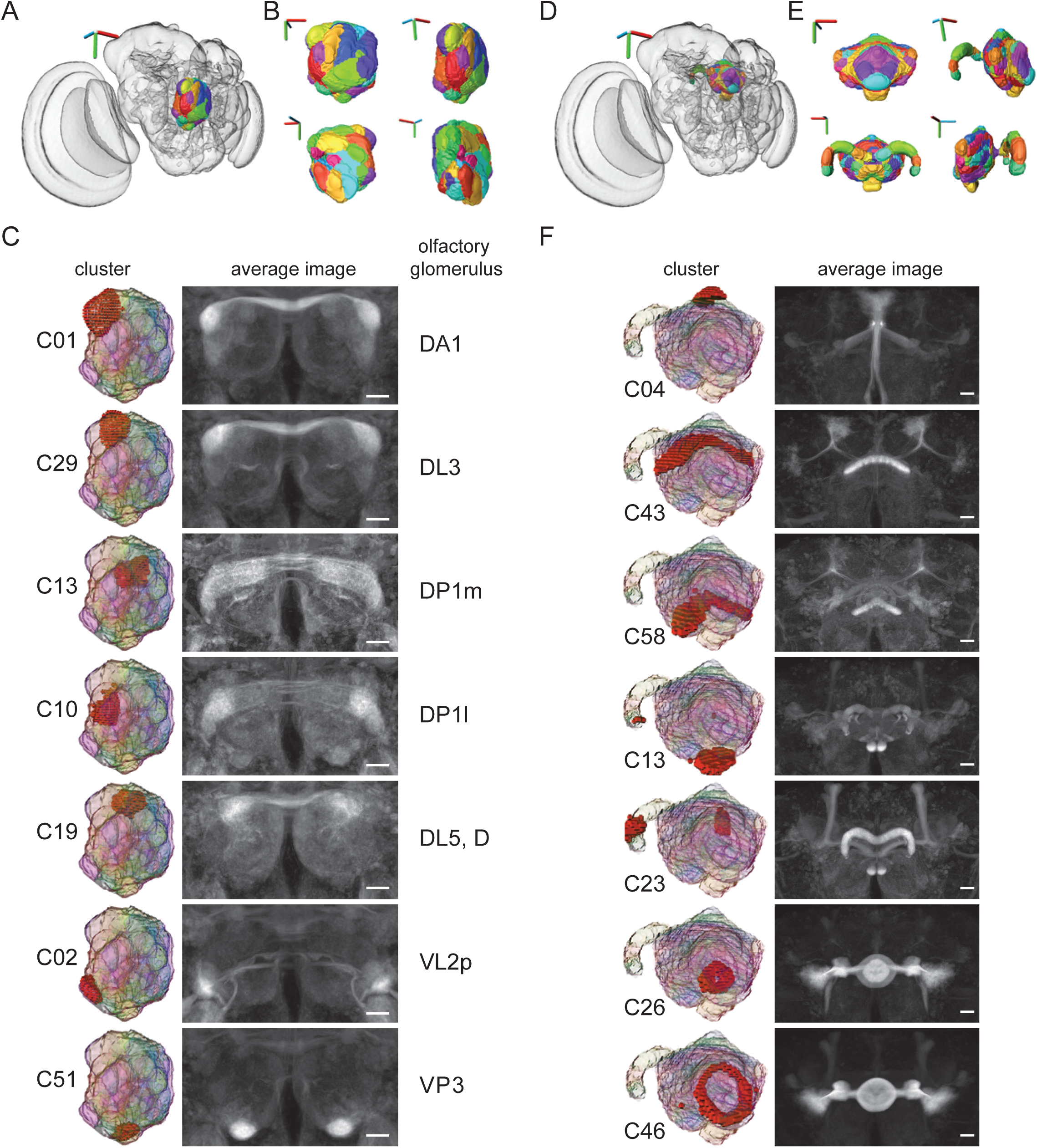
Automatic segmentation of antennal lobe (AL) and central complex (CX). A) The automatic clustering results from the right AL plotted in the whole brain. 3D axes scale 40 μm. B) 3D views of the AL clustering assignments. 3D axes scale 15 μm C) individual clusters (left), average image of strongly expressing driver lines with broad driver lines removed (middle), and manually assigned corresponding olfactory glomerulus (right). Scale bars 20 μm. D) The automatic clustering results from CX plotted in the whole brain. 3D axes scale 40 μm. E) 3D views of the CX clustering assignments. 3D axes scale 30 μm. F) individual clusters (left), average image of strongly expressing driver lines with broad driver lines removed (right). Scale bars 20 μm. (Panels A-C: Janelia FlyLight data for the right AL, run 1, 23769 voxels, 3462 driver lines, *k* equal 60. Panels D-F: Janelia FlyLight data for CX, run 1, 27598 voxels, 3462 driver lines, *k* equal 60.)

Individual clusters were highlighted (Figure 2C, left column) and used to look at the individual GAL4 lines that have particularly high expression within a given cluster (see https://strawlab.org/braincode) or to take an average of all confocal image stacks from all GAL4 lines that strongly present in a particular cluster but not broadly expressing elsewhere in the target brain region (Figure 2C, right column, Figure 2–figure supplement 2,3). Although our input brain region was the right AL, the average image stacks show a high level of symmetry across the midline. Furthermore, a large fraction of voxels belonging to a given glomerulus whose identity was manually assigned in an nc82 stained brain as ‘ground truth’ were shared with individual clusters (Figure 2-figure supplement 1). In a subsequent manual step, we used these correspondences to identify automatically extracted clusters as specific olfactory glomeruli (Figure 2C). When the same analysis was performed on an entirely independent dataset (from the Vienna Tiles collection rather than the Janelia FlyLight) the results were qualitatively similar (Supplementary file 1 and https://strawlab.org/braincode website).

### Central complex, Mushroom bodies, Sub-esophageal zone

We performed further clustering on both relatively well-understood brain regions and the ‘terra incognita’ of diffuse neuropils. The central complex (CX) has been the focus of substantial anatomical work (Bausenwein et al., 1986; Hanesch et al., 1989; Lin et al., 2013; Strauss and Heisenberg, 1993) and has been recently described in extensive detail using split-GAL4 line generation and manual annotation (Wolff et al., 2015). The Braincode algorithm automatically identified many of the prominent structures within this brain region (Figure 2D-E). For example, individual shells of the ellipsoid body neurons are segmented, individual layers of the fan shaped body are found, and the protocerebral bridge is segmented into distinct regions. In this case, our input brain region spanned the midline to cover the entire CX region, and consistent with expectations for a working algorithm, the clustering results are mirror symmetric across the midline (Figure 2F, Figure 2–figure supplement 4,5).

The results on these well studied brain regions therefore support the idea that patterns of coexpression can indeed be used to identify functional units and that the Braincode algorithm is capable of automatically segmenting brain regions into putative, biologically meaningful sub-regions.

On the https://strawlab.org/braincode website, we also include the results of clustering the mushroom bodies (MBs) and sub-esophageal zone (SEZ). Future clustering results can be added upon request.

### Optic glomeruli

The posterior ventrolateral protocerebrum (PVLP), posterior lateral protocerebrum (PLP) and anterior optic tubercle (AOTU) are diffuse neuropils to which the majority of outputs from the medulla and lobula neuropils within the optic lobes project (Otsuna and Ito, 2006; Strausfeld and Bacon, 1983; Strausfeld and Lee, 1991). By analogy to the antennal lobes, where a single glomerulus processes the output of a single type of olfactory sensory neuron (OSN), it is proposed that a single VPN type projects to a single optic glomerulus and encodes a single visual feature (Mu et al., 2012). These regions have accordingly received some attention, but the specific location and identity of structures within these regions remains incompletely described. Therefore, we used Braincode to identify putative functional units in this region (Figure 3AB). We call the union of these three neuropils (PVLP, PLP and AOTU) the optic Ventrolateral Neuropil (oVLNP).

**Figure 3.**
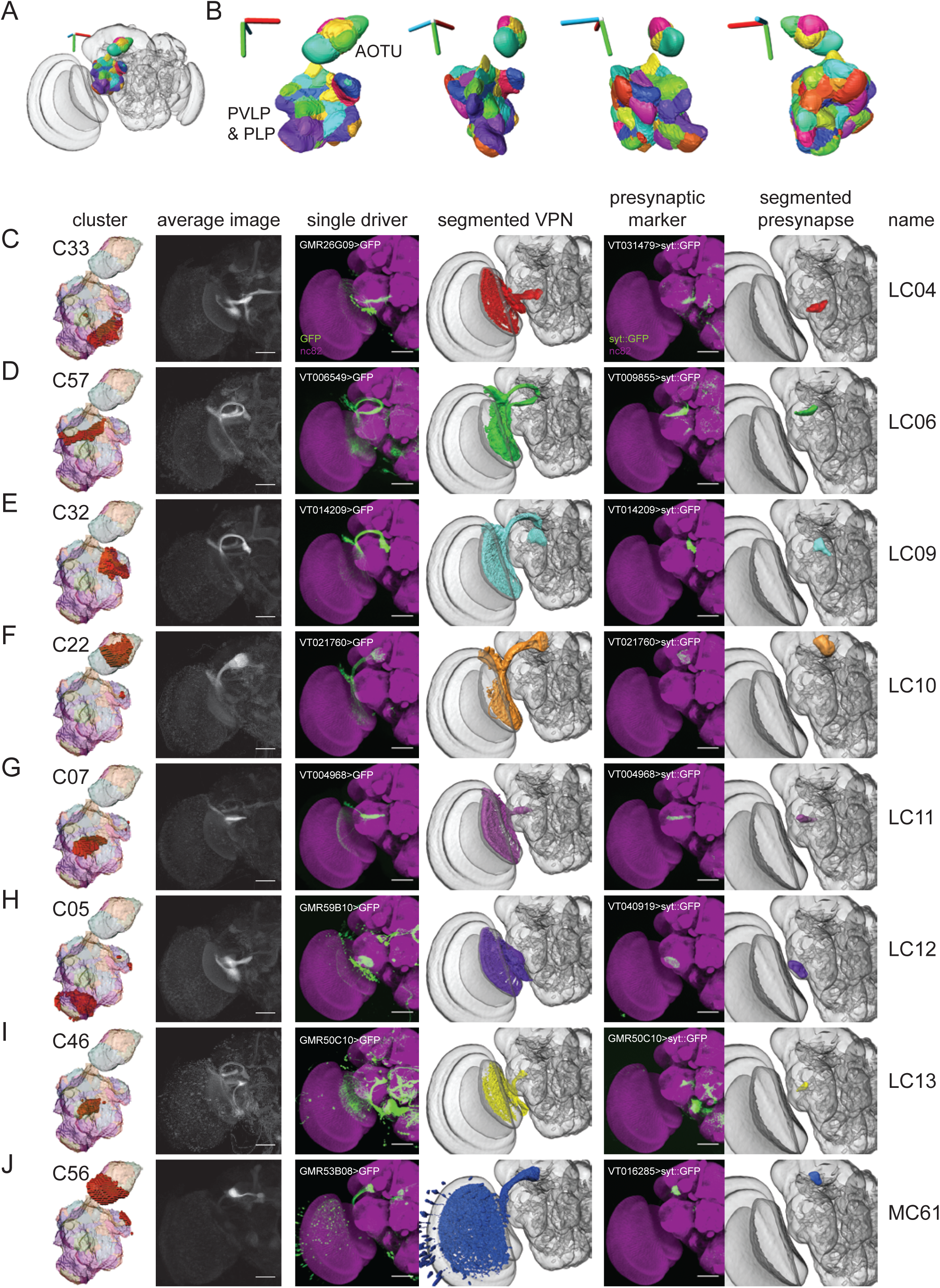
Automatic segmentation reveals clusters that correspond to optic glomeruli associated with previously identified visual projection neurons (VPNs). A) Clusters from the oVLNP region plotted within entire brain. 3D axes scale 40 μm. B) Multiple 3D views of clusters. 3D axes scale 40 μm. C-J) Individual clusters, average images, selected driver lines, 3D segmentations of a particular VPN type, presynaptic marker (UAS-synaptotagmin::GFP) expressed by a single driver and 3D segmentation of presynaptic region to define optic glomerulus. (All panels: Janelia FlyLight data for the oVLNP region defined as PLP, PVLP, and AOTU, run 1, 42317 voxels, 3462 driver lines, *k* equal 60. Scale bars 50 μm.)

Consistent with the idea that some of the automatically segmented clusters are optic glomeruli, we could identify a single, previously described VPN type projecting to many of these clusters (Figure 3C-J). In addition to creating an average image by combining driver lines expressing in the cluster, we selected individual driver lines that appeared to drive expression in a single VPN type projecting to this cluster. By comparing the morphology of the neurons selected this way with previous reports, particularly Otsuna and Ito (2006), we could identify LC04, LC06, LC09, LC10, LC11, LC12, LC13 and LC14. (Missing elements from the sequence – LC01, LC02, LC03, LC05, LC07 and LC08 – were omitted by Otsuna and Ito due to uncertain identification compared to previous work.) To image the precise location of synaptic outputs of each of these VPN types, we expressed a presynaptic marker, synaptotagmin::GFP (syt::GFP) (Zhang et al., 2002), using the selected driver lines. After registering these newly acquired confocal image z-stacks to the templates of the Vienna or Janelia collections, we could then define the 3D location and extent of the VPN output – the VPN’s associated optic glomerulus – by performing assisted 3D segmentations of the presynaptic regions. Initial inspection showed a substantial similarity between such manually validated optic glomeruli and automatically identified clusters, and below we quantify this correspondence.

When segmenting a large brain region into putative functional units, we might expect to find axon tracts in addition to nuclei or glomeruli. Indeed, the clustering results also included two apparent axon tracts through this region, the great commissure connecting the two contralateral lobulae including LC14 and the tract that includes the Lat (lamina tangential) neuron type (Figure 4).

**Figure 4.**
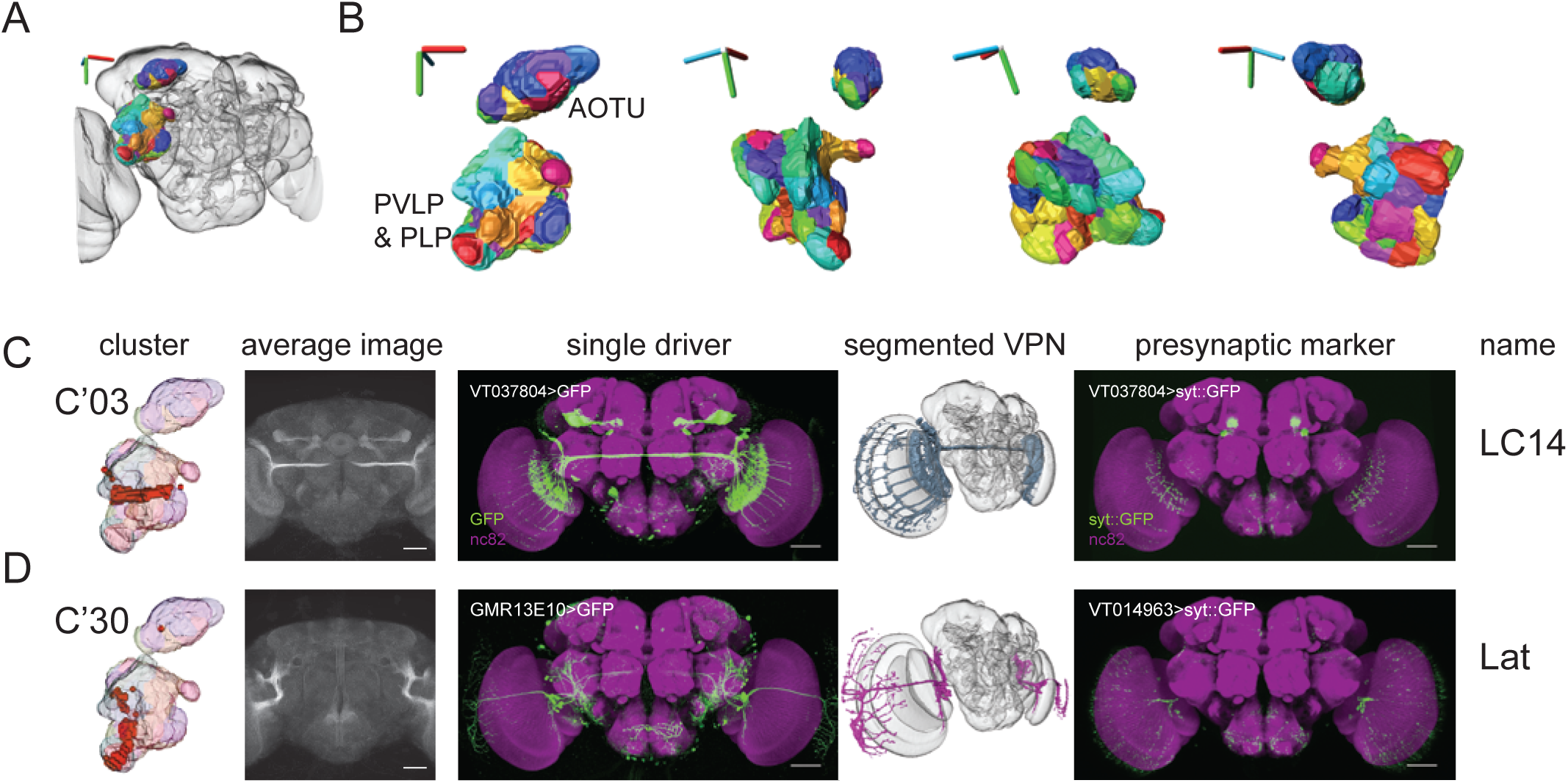
Automatic segmentation reveals clusters that correspond to tracts associated with previously identified visual projection neurons. A) Clusters of the oVLNP with the Vienna Tile dataset plotted within entire brain. 3D axes scale 40 μm. B) Multiple 3D views of clusters. 3D axes scale 30 μm. C) Cluster associated with the giant commissure, including LC14 neurons. D) Cluster associated with the axons of Lat neurons. (All panels: Vienna Tiles data for the oVLNP, run 1, 13458 voxels, 6022 driver lines, *k* equal 60. Scale bars 50 μm.)

In addition to clusters corresponding to output regions of previously identified neuron types, we found clusters that appear to be projection targets of VPNs that have not been previously described. These novel VPNs are eight lobula columnar (LC) types, four lobula plate-lobula columnar (LPLC) types, one lobula-plate columnar type, and two medulla columnar (MC) VPNs types. Using the same presynaptic GFP expression approach as above, we saw substantial similarity between these manually validated optic glomeruli to the clustering result (Figure 5,6). For each cell type, we used the FlyCircuit database (Chiang et al., 2011) to identify multiple example single neuron morphologies (Figure 8-table supplement 1). We named these neuron types by continuing the sequence onwards from the last published number for a particular class (i.e. LC15 is the first lobula columnar type we identified whereas LC14 was previously reported).

**Figure 5.**
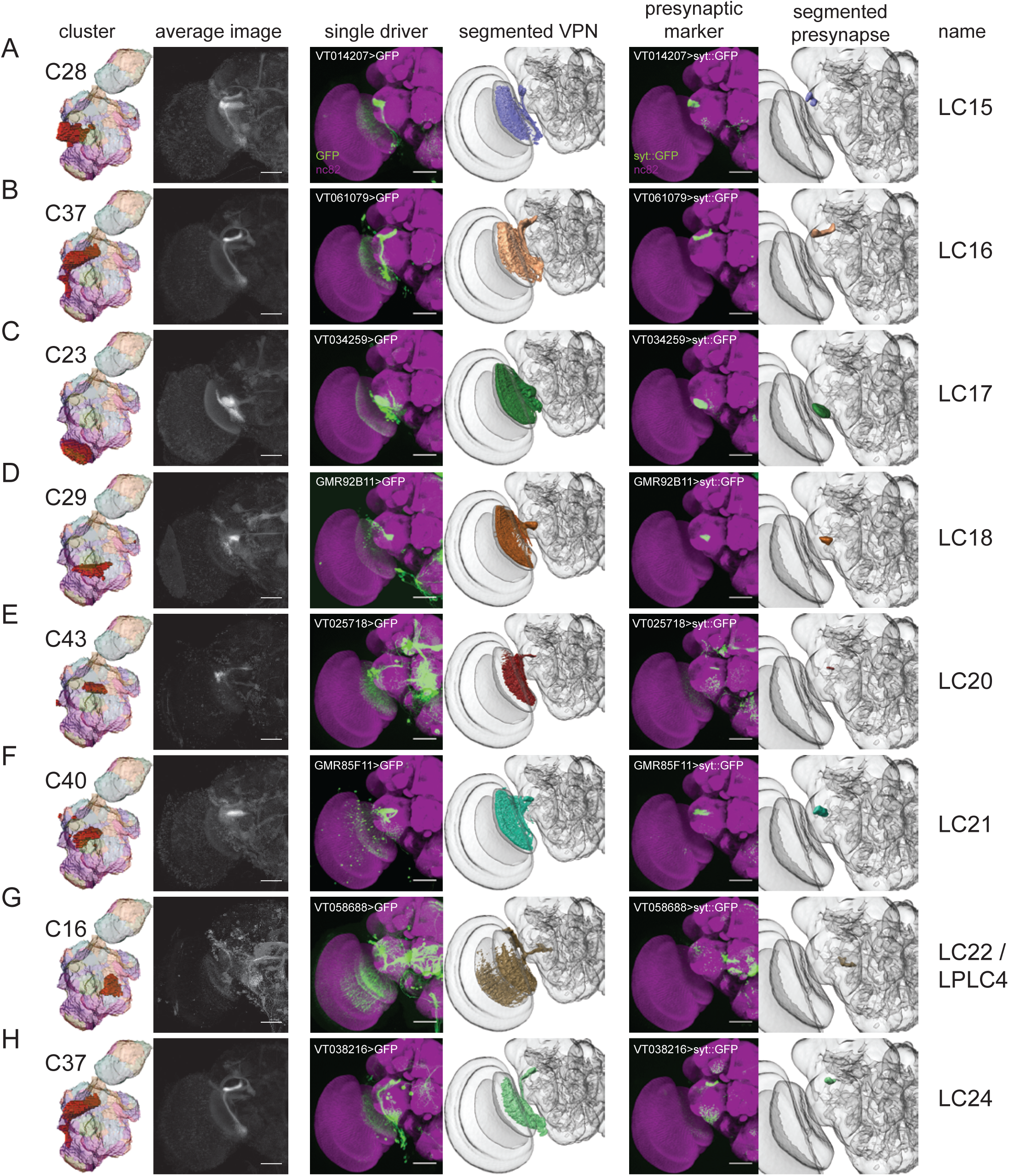
Automatic segmentation reveals clusters that correspond to optic glomeruli associated with newly identified LC-type visual projection neurons. A-H) Individual clusters, average images, selected driver lines, 3D segmentations of a particular VPN type, presynaptic marker (UAS-synaptotagmin::GFP) expressed by a single driver and 3D segmentation of presynaptic region to define optic glomerulus. (All panels: Janelia FlyLight data for the oVLNP, run 1, 42317 voxels, 3462 driver lines, *k* equal 60. Scale bars 50 μm.)

**Figure 6.**
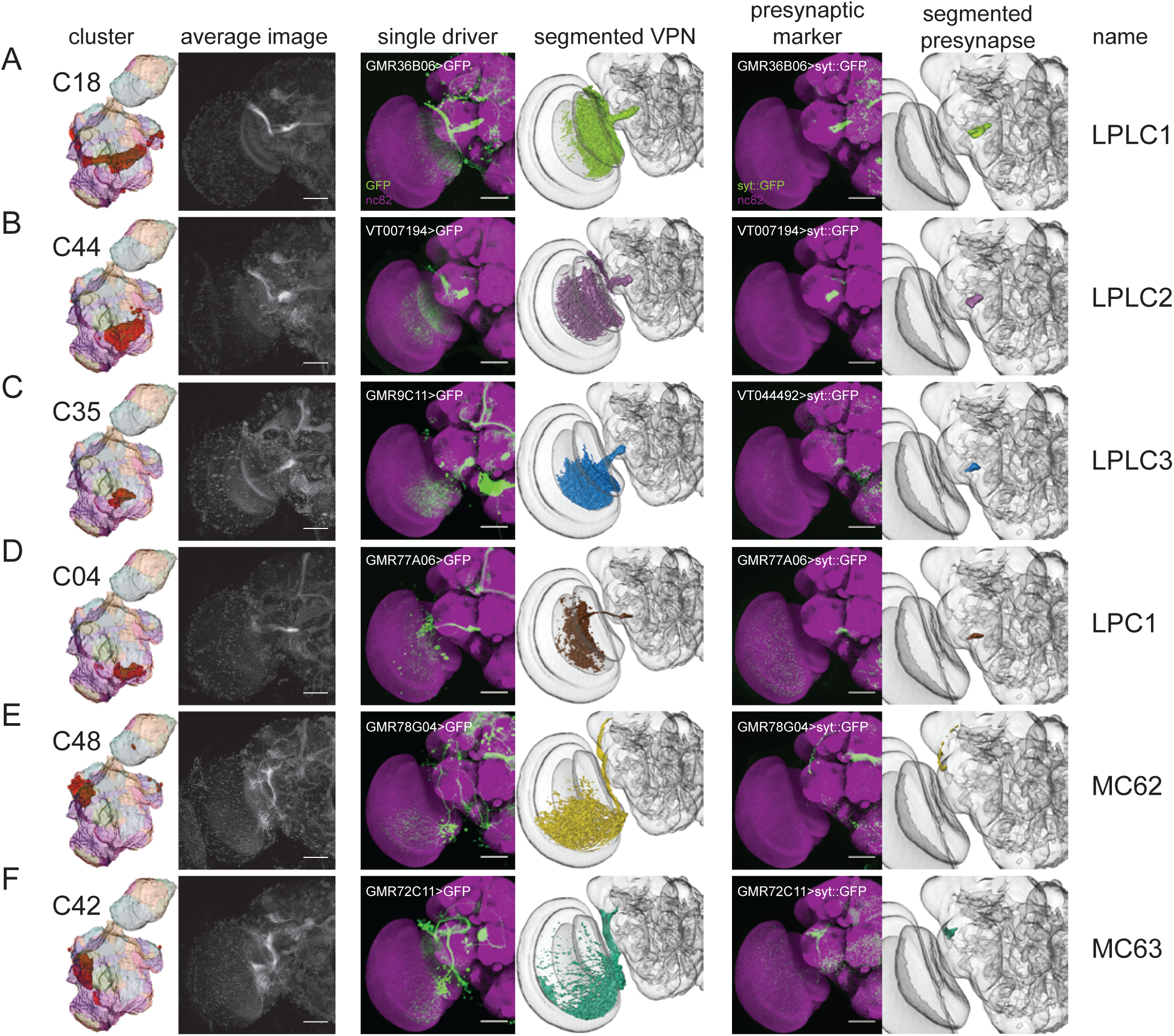
Automatic segmentation reveals clusters that correspond to optic glomeruli associated with newly identified LPLC, LPC, and MC-type visual projection neurons. A-F) Individual clusters, average images, selected driver lines, 3D segmentations of a particular VPN type, presynaptic marker (UAS-synaptotagmin::GFP) expressed by a single driver and 3D segmentation of presynaptic region to define optic glomerulus. (All panels: Janelia FlyLight data for the oVLNP, run 1, 42317 voxels, 3462 driver lines, *k* equal 60. Scale bars 50 μm.)

We defined the precise 3D location of the optic glomeruli by segmenting the presynaptic marker signal from registered confocal image stacks of VPN lines. Quantification showed a high degree of colocalization between these manually validated optic glomeruli and voxels from specific clusters, and plotting these results showed that the Braincode method automatically produces segmentations with substantial similarity to those derived from labor-intensive manual techniques (Figure 7A). This holds true across a second, entirely distinct dataset (Figure 7B).

**Figure 7.**
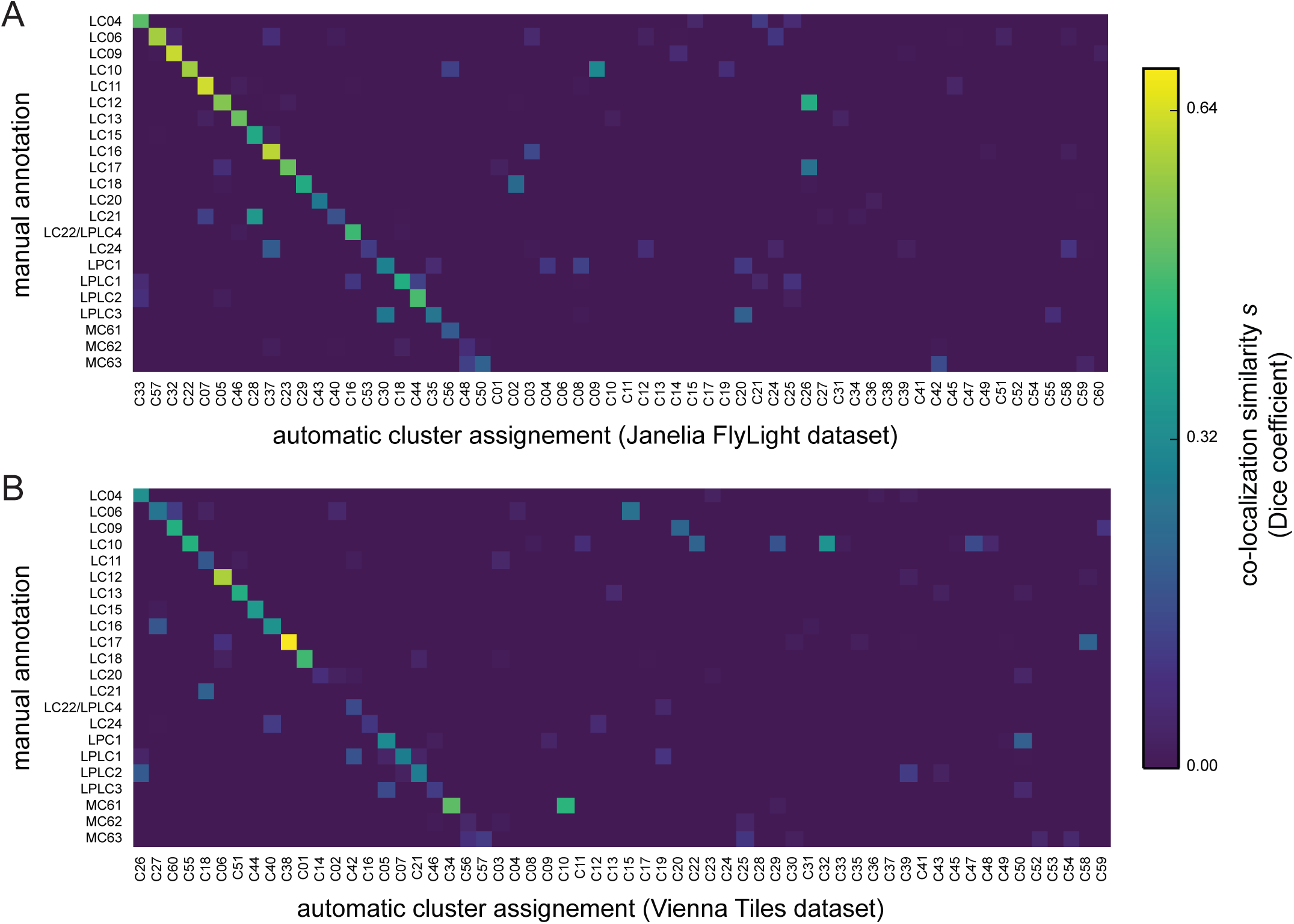
Automatically assigned clusters colocalize with manually segmented optic glomeruli. A) Colocalization similarity (measured based on set of voxels in manually annotated region and set of voxels in clustering result) between the Janelia FlyLight dataset and manual assignments using the same 3D template brain. (Janelia FlyLight data for run 1, oVLNP, 42317 voxels, 3462 driver lines, *k* equal 60.) B) Colocalization similarity between the Vienna Tiles dataset and manual assignments using the same 3D template brain. (Vienna Tiles data for run 1, oVLNP, 13458 voxels, 6022 driver lines, *k* equal 60.)

We evaluated completeness of the results in two ways. First, we clustered both data sets twice with *k* equal 60 but different random number seeds and discovered in each run at least 23 of the 25 glomeruli or tracts associated with a particular VPN type (Figure 8–table supplement 1). We expect subsequent repetitions to reveal few, if any, additional novel structures. Secondly, we noted that regions of high intensity anti-Bruchpilot (nc82 antibody) staining, an indicator of synaptic contacts, coincide with optic glomeruli. In the brain regions investigated, we found glomeruli for all such high intensity regions (Figure 8). We did not perform clustering on the Posterior Slope (PS), a region targeted by the lobula plate tangential cells (LPTCs), and thus did not expect to find any clusters associated with these neurons, nor did we find any such clusters. Taking these results together, we conclude that the Braincode method can find a majority of structures in a particular region.

**Figure 8.**
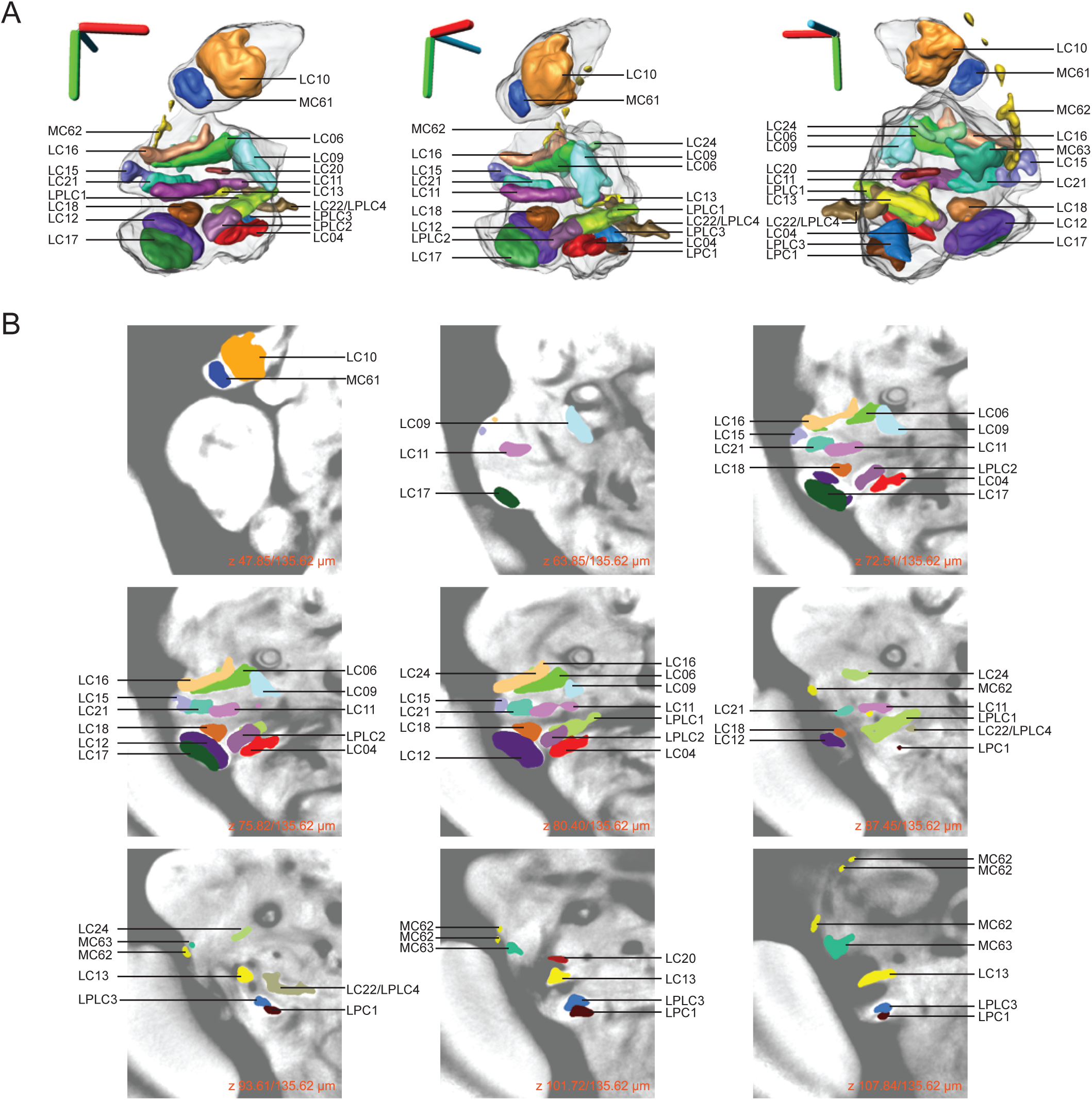
An atlas of the optic glomeruli defined by manual segmentation of presynaptic marker expression experiments. A) 3D rendering of all identified optic glomeruli registered onto a 3D reference brain. Optic glomeruli were segmented from single driver confocal images expressing presynaptic marker (UAS-synaptotagmin::GFP). (Scale bars 40 μm.) B) Z-stack showing the location of each optic glomerulus in a 2D view on the background of an average image of many individual nc82 stained brains.

### Interpreting results from automatic clustering

As noted above, any clustering algorithm has a parameter that (implicitly or explicitly) controls the number of resulting clusters. An important question when using these algorithms, then, is how to set that parameter. In the ideal case, an inherent clustering is easy to identify within the data and nearly trivial for an automatic algorithm to extract. Often however, and we believe this is the case for the type of spatial expression data used here, the distinctions between different portions of the data are somewhat unclear and the clustering algorithm creates a classification which may be different from an expert assessment. Experts themselves often disagree, however, due to debates in which ‘lumpers’ argue that differences are insignificant and only obscure a more important deeper unity and ‘splitters’ argue that the differences seen reflect important underlying distinctions. Therefore, we expected some degree of splitting, lumping or both in our results.

To evaluate the distinctness of our clusters and to gain insight into the molecular distances between different clusters, we plotted distance matrices between medoids (Figure 7–figure supplement 1 A,C). We also made use of t-distributed stochastic neighbor embedding (von der Maaten and Hinton, 2008) to make 2D plots in which medoids are plotted in close proximity when their molecular distance is low but farther apart when they are less closely related (Figure 7–figure supplement 1 B,D). In some cases, this approach shows that some clusters identified as distinct have a small ‘molecular distance’ and thus might be considered to result from excessive splitting. On the other hand, evidence of potential lumping comes from cases such as only a single cluster being found for the optic glomeruli corresponding to the LC16 and LC24 VPN types, despite the fact that manual segmentations of their associated optic glomeruli showed that these project to anatomically distinct (but adjacent) regions (Figure 5B,H). Despite a potentially unsolvable assignment problem of the existence one or two ‘true’ functional units, co-clustering indicates that there are some driver lines that drive expression in both glomeruli.

One illustrative example of the challenge of whether to lump and split comes from the optic glomerulus associated with the LC10 neuron type. Clusters C09 and C22 in run 1 of the Janelia Fly Light dataset (Figure 3-figure supplement 1) correspond to dorsal and ventral parts of the medial AOTU respectively, and the LC10 neuron type projects to both clusters. While LC10 subtypes – with distinct morphology and with inputs from distinct layers of the lobula – have been identified that target these regions preferentially (Costa et al., 2015; Otsuna and Ito, 2006), our results – separate clusters but very low distance on the t-distributed stochastic neighbor embedding (t-SNE) plot (Figure 7-figure supplement 1 B) – suggest that there is relatively little molecular distance between the dorsal and ventral parts of the medial AOTU. Indeed, after searching through the list of driver lines with substantial expression in C22, we could find only a single driver line, GMR22A07-GAL4, that drove strong expression in a VPN targeting this region and had specificity for Otsuna and Ito’s (2006) LC10a subtype but not LC10b. It would be tempting to conclude, then, that the division of the medial AOTU was erroneously split by the clustering algorithm. Yet the existence of distinct LC10 subtypes suggests that there are genuine, if small, distinctions between these regions. We suggest that the LC10 neuron type presents an example of the lumping versus splitting problem within spatial expression data. It may be that further data, for example detailed studies on LC10 subtype morphology and molecular expression, could resolve the issue. In the absence of such data, subdividing large brain regions can be useful simply as a way to reduce the complexity of a large brain region and need necessarily imply a strong claim of correspondences to genuine anatomical correlates. And this benefit of clustering would furthermore remain even if further data did not support a clear conclusion.

As discussed, automatic calculation of a measure of repeatability (adjusted Rand index, Figure 1–figure supplement 1) found no obvious optimum value of *k*. Therefore, we sought to gain a more biologically meaningful sense of consistency across multiple runs of the algorithm for the value of *k=60* that we chose by performing a visualization comparing the results of a manual segmentation of a brain region with the automatic segmentations. We did this for the oVLNP with each of four different clustering runs, two from each dataset (Figure 7A,B and Figure 7-figure supplement 2A,B). The results show that, despite different random number initialization seeds, most optic glomeruli have a strong correspondence with a single cluster across repeated runs of the algorithm within and across the two datasets (Vienna Tiles and Janelia FlyLight). This suggests substantial biologically meaningful repeatability within and between datasets.

In sum, we suggest that the automatic segmentations produced by Braincode should be used as hypotheses that must be further investigated, as we have done here for the visual system, before strong conclusions can be drawn about intrinsic neuroanatomical structure.

### Little VPN convergence to single optic glomeruli

Of the 22 optic glomeruli we identified, only a single one was targeted by two VPN types. Apart from LC22 and LPLC4 projecting to the same glomerulus, we found no other instance of convergence of multiple VPN types to a single optic glomerulus. In some cases however, two VPN types projected to a single cluster. For example, LC11 and LC21 both project to the region containing C07 (Figure 7). While there are some regions of presynaptic colocalization in the underlying signals in registered images, there are also non-overlapping presynaptic localizations and thus the data suggest that the glomeruli are at least partially distinct (Figure 8B). LC12 and LC17 are another similar pair but the presynaptic localization is even more distinct in this case (Figure 8B). Similarly, the presynaptic localizations of LC16 and LC24 both are within cluster C37, although in this case we think that a paucity of driver lines driving expression in LC24 likely precluded a separate cluster from being identified. In summary, with a single exception, we do not find evidence for multiple VPNs projecting to a single optic glomerulus and instead propose that where we do see projection to the same cluster that this results from lumping within the clustering algorithm.While we cannot exclude the possibility that more optic glomeruli exist that are the targets of two or more VPN types, our data show that such cases are exceptional. Conversely, we found that each VPN type projects to a single glomerulus. Together, these two observations allow us to propose naming optic glomeruli according to the VPN type(s) that project to them.

### A map of the optic glomeruli of *Drosophila*

We can synthesize the novel findings of this automatic and manual characterization of this brain region with a movie showing segmented visual projection neurons and the presynaptic output regions associated with each of these VPNs (Video 1). Furthermore, we have created reference figures describing the optic glomeruli as the targets of specific VPNs (Figure 8) and provide separate 3D models of each VPN type and its associated optic glomerulus all in a common 3D template brain coordinate system (Supplementary file 1).

**Video 1.**
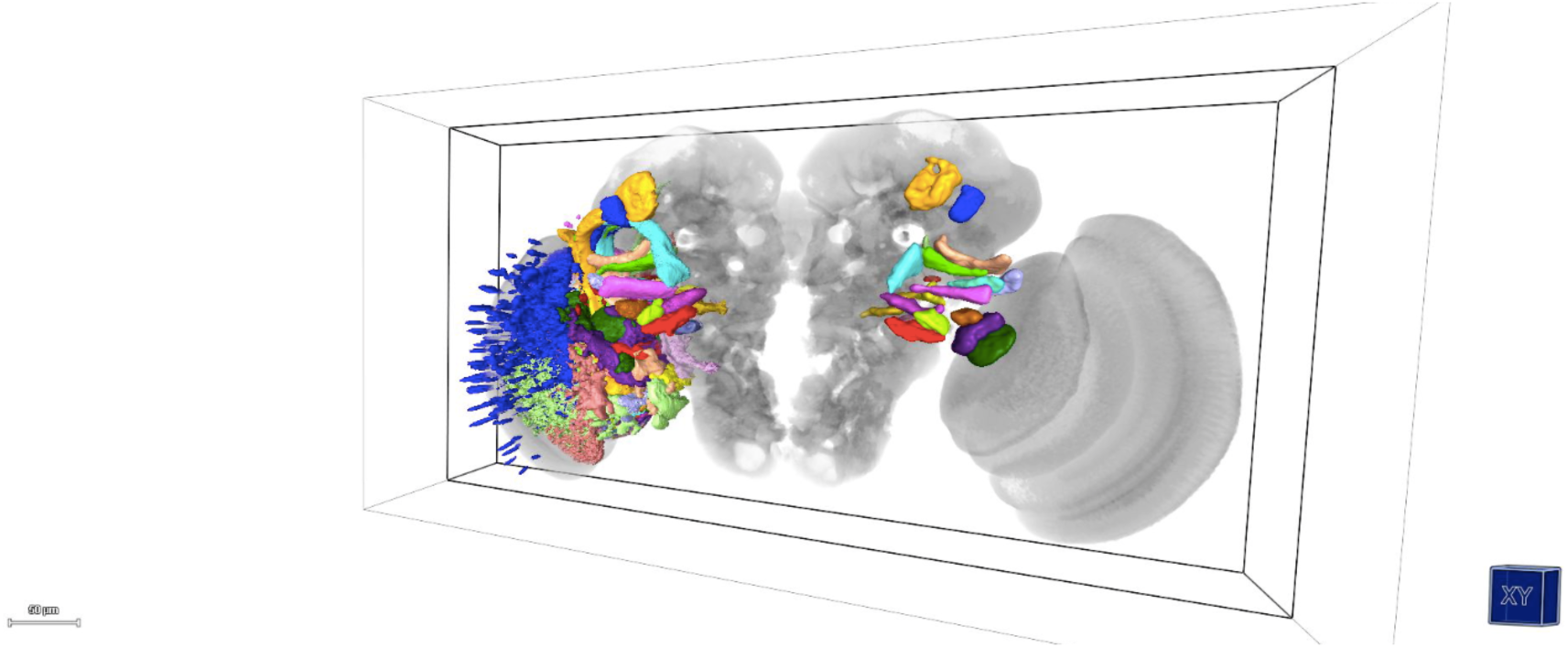
3D location of manually segmented visual projection neurons and optic glomeruli. Right half shows 3D rendering of all identified optic glomeruli registered onto a 3D reference brain. Optic glomeruli were segmented from single driver confocal images expressing presynaptic marker (UAS-synaptotagmin::GFP). Left half shows 3D rendering of visual projection neurons segmented from single driver confocal images expressing a non-localized cell membrane marker (UAS-CD8::GFP).

### Pathways leaving the optic glomeruli

Just as we identified driver lines expressing in VPN types that enter a particular optic glomerulus, we can also use the lists of driver lines expressed in a given cluster to suggest candidate interneurons that are largely contained within a particular glomerulus or projection neurons that leave from the glomerulus. To demonstrate the potential of this approach, we used such driver lines to drive expression of two reporters, a red fluorescent dendritic marker UAS-DenMark::mCherry (Nicolaï et al., 2010) and a green fluorescent presynaptic marker UAS-Syt::GFP (Zhang et al., 2002). In several cases, we can identify candidate neurons that appear to have dendritic inputs in a particular glomerulus and project elsewhere in the brain (Figure 9).

**Figure 9.**
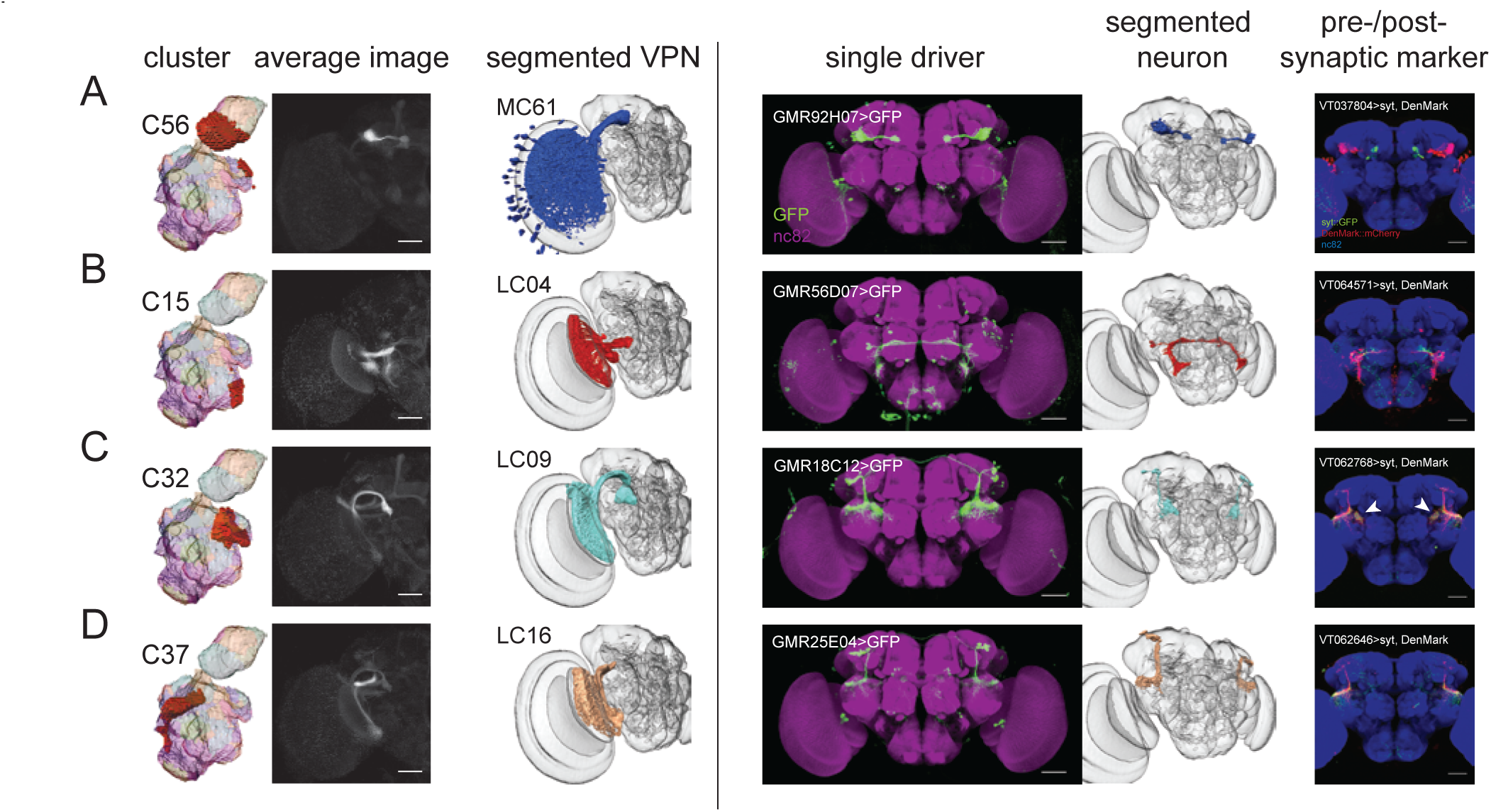
Using clusters to identify neuron types that express dendritic markers in a particular optic glomerulus and project to another region. A-D) Neurons that project to (left) and from (right) a particular optic glomerulus, found using candidate searches from the Braincode result lists. Pre- and post-synaptic markers were UAS-synaptotagmin::GFP and UAS-DenMark::mCherry, respectively. A) Putative outputs from the optic glomerulus to which MC61 projects include a neuron type that projects to the bulb. Such cells express post-synaptic marker in the AOTU and pre-synaptic markers in the bulb. (Driver lines: GMRH07-GAL4, VT037804-GAL4) B) The optic glomerulus to which the LC04 neuron type projects contains a neuron, likely the giant commissural interneuron CGI (Phelan et al., 1996) that expresses post-synaptic marker in this glomerulus. (Driver lines: GMR56D07-GAL4, VT064571-GAL4) C) The optic glomerulus to which the LC09 neuron type projects contains a neuron that expresses pre- and post-synaptic markers in this glomerulus (arrowheads). (Driver lines: GMR18C12-GAL4, VT062768-GAL4) D) The optic glomerulus to which the LC16 neuron type projects contains a neuron that expresses pre- and post-synaptic markers in this glomerulus. (Driver lines: GMR25E04-GAL4, VT062646-GAL4)

## Discussion

We have demonstrated that applying a clustering algorithm to imaging data from large-scale enhancer libraries segments brain regions into smaller, putative functional units such as glomeruli and axon tracts. When applied to *Drosophila* data, automatically extracted clusters have a high correspondence with glomeruli and other neuropil subdivisions within the antennal lobes and central complex, suggesting the utility of the approach. We used this approach to inform a detailed investigation of the optic Ventrolateral Neuropil (oVLNP), a region where most outputs from the medulla and lobula neuropils within the optic lobes reach the central brain. We identified several neuron types that, to the best of our knowledge, have not been previously described: eight lobula columnar (LC) neuron types, four lobula plate-lobula columnar (LPLC) types, one lobula-plate columnar type, and two medulla columnar (MC) types.

We found a nearly one-to-one projection of visual projection neurons to optic glomeruli. This is consistent with the idea that each optic glomerulus processes input from a single cell type and is therefore similar to the olfactory glomeruli in the sense that a dedicated glomerulus receives input from a single distinct input cell type (Mu et al., 2012). Future work could investigate whether the regions are homologous in an evolutionary sense and if the similarities extend to functional aspects and developmental mechanisms.

Recent computational neuroanatomical work has sought to use extensive collections of registered image stacks from stochastically labeled brains (Chiang et al., 2011) to identify cell types (Costa et al., 2015) construct a mesoscale connectome of the fly brain (Shih et al., 2015) or to find groups of morphologically similar neurons likely from the same neuroblast (Masse et al., 2012). Given the complementary strengths of the respective approaches – resolution to the single-cell level with stochastic labeling approaches and candidate driver lines and molecular identity from the Braincode approach, it may be productive to perform further analysis that takes advantage of these differences. For example, it might be possible to perform a motif analysis to identify enhancer fragments correlating with anatomical features such as projection target, axon tract location, or branching pattern. Additionally, because the enhancer fragments are likely to regulate genes that neighbor the enhancer region in the genome (Kvon et al., 2014), this approach could be used to suggest genes that are particularly distinct for specific brain regions and potentially for specific cell types.

The approach outlined here has several technical dependencies, which may represent limitations in some cases. Firstly, there is an obvious requirement that any structure segmented automatically must have a physical scale at least comparable to, if not larger than, the error in registering multiple samples. Secondly, enough registered enhancer line images must be available to provide a signal sufficient for clustering. Third, underlying biological variability in the developmental patterns must be less than the variability in the registered expression data. In addition to these technical dependencies, we found that the use of an automatic classification algorithm does not solve the classic ‘lumper versus splitter’ problem. Also, while we have shown that clustering often identifies regions with anatomical correlates such as a glomerulus, in other cases this may be less clear. In any case, the clusters identified result from patterns of expression in many driver lines but it may be that only some driver lines are confined to the boundaries of a given cluster. In cases where the automatically extracted clusters do not clearly correspond with an anatomical structure, we propose that clustering may nonetheless be useful in reducing the complexity of thinking about a large brain region by dividing it into smaller elements.

Despite these potential limitations, the Braincode approach is not limited to *Drosophila*. Data are available from recent Zebrafish enhancer trap experiments (Kawakami et al., 2010; Kondrychyn et al., 2011) and registering brains is also possible (Ronneberger et al., 2012). Together, these would enable an attempt to apply the Braincode technique. New developments, such as the use of site-specific integrase (Lister, 2011; Mosimann et al., 2013) could be used to minimize expression level variation due to effects of where a transgene integrates in the genome and improve efficiency and thus produce comparable datasets to those used here for *Drosophila*. Such an effort in Zebrafish could be used to suggest driver lines corresponding to functional units identified in brain-wide activity-based experiments (Ahrens et al., 2012; Kubo et al., 2014; Portugues et al., 2014; Randlett et al., 2015). Similar datasets are being gathered in another fish species, Medaka (Alonso-Barba et al., 2015). Variability of brain development in mammals may make the approach more challenging, or only operate on larger scales, in these species. Nevertheless, the ability to automatically segment brain regions into putative functional units could prove useful in unraveling structure-function relationships in a variety of species.

## Methods and materials

### Drosophila Strains/Stocks

Flies were raised at 25 degrees Celsius under a 12 hour light-dark cycle on standard cornmeal food. Used GAL4 lines were from the Vienna Tiles collection (generated by the groups of B.J. Dickson and A. Stark, unpublished data, see also Kvon et al., 2014) and Janelia GAL4 library (Pfeiffer et al., 2010, 2008) and were obtained from the Vienna Drosophila RNAi Center or Bloomington Drosophila Stock Center (BDSC), respectively. UAS-mCD8::GFP was generated by B.J. Dickson group. UAS-DenMark::mCherry, UAS-synaptotagmin::GFP was created by B.A. Hassan and obtained from BDSC.

### Sample Preparation and Imaging

Fly dissection and staining were performed as previously described (Yu et al., 2010) using 3 to 5 days old adult flies. In brief, brains were dissected in phosphate buffered saline (PBS), fixed in 4 % paraformaldehyde in PBS with 0.1 % Trition-X-100 and subsequently blocked in 10 % normal goat serum (Gibco Life Technologies). Brains were incubated in primary and secondary antibodies for a minimum of 20 hours at 4 degrees Celsius and washed in PBS with 0.3 % Trition-X-100. Fly brains were mounted in Vectashield (Vector Laboratories). We used the following primary antibodies: rabbit polyclonal anti-GFP (1:5000, TP401, Torrey Pines), mouse monoclonal anti-bruchpilot (1:20, nc82, Developmental Studies Hybridoma Bank), chicken polyclonal anti-GFP (1:10.000, ab13970, Abcam), rabbit polyclonal anti-DsRed (1:1000, 632496, Clontech). We used the following secondary antibodies: Alexa Fluor 488, 568 or 633 antibodies (1:500 to 1:1000, Invitrogen Life Technologies).

Images were acquired using point scanning confocal microscope LSM780 or LSM700 (Zeiss) equipped with 25x/0.8 plan-apochromat multiimmersion or 20x/0.8 plan-apochromat dry objectives, respectively. To avoid channel crosstalk confocal Z-stacks were recorded in the multi-track (LSM700) or online fingerprinting mode (LSM780).

### Registration, Assisted Segmentation, and 3D-Rendering

For both datasets an intensity-based nonlinear warping method was used. For the Vienna Tiles dataset we used the approach described in (Yu et al., 2010) and for the Janelia dataset, brains were registered according to (Cachero et al., 2010). Fiji (ImageJ) and Amira (4.1.2, Mercury Computer Systems) software were used for image processing and analysis. Amira label field function was used to segment optic glomeruli, projections and neuron types from registered images. Surface files of segmented structures were generated using constrained smoothing for full neuron segmentations and unconstrained smoothing for optic glomeruli. We additionally used the BrainGazer visualization software (Bruckner et al., 2009). In all 3D figures, we included a 3D axes scale in which red specifies the lateral axis with positive towards the animal’s left side, green specifies the dorsal-ventral axis with positive towards ventral, and blue specifies the anterior-posterior with position towards posterior. Due to the use of a perspective projection in these figures, the size of the 3D axes scale is only approximate.

### Thresholding, Dice similarity, k-Medoids, and t-SNE

GAL4 expression patterns were transformed into a binary representation in two steps. First, the image is thresholded and second, morphological opening with a 3×3×3 kernel is applied to reduce clutter. The threshold was chosen so that the resulting mask yielded 1% stained voxels. This simple heuristic was more reliable for the datasets tested compared to other standard automatic thresholding methods.

From the binarized images, the set of expressing lines was assembled for each voxel. Similarity between voxels based on the respective expression set from voxel A and the set from voxel B is computed using Dice’s coefficient as 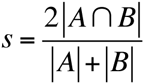 where ∩ denotes intersection and *|*χ| denotes the number of elements in set χ. To decrease the effects of registration error and image acquisition noise and to increase the speed of subsequent processing steps, we binned the original image voxel data into larger voxels, typically a 3×3×3 downsampling. The k-medoids algorithm (Kaufman and Rousseeuw, 1987) was run in Julia 0.4.0 using JuliaStats Clustering 0.5.0 (see Supplementary file 1). The *k*-medoids was performed on Dice dissimilarity (1-*s*). To visualize the distance between medoids, we used the implementation of t-distributed stochastic neighbor embedding (von der Maaten and Hinton, 2008) in Python 2.7.10 using the Scikits Learn 0.16.1 software package (Pedregosa et al., 2011) with precomputed distances using metric distance 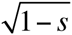 between medoids.

### Nomenclature

Existing nomenclature was used for previously identified neuron types when an unambiguous match was possible. Lobula columnar neurons were first systematically described in *Drosophila* in (Fischbach and Dittrich, 1989) which called these ‘Lcn’ types and included Lcn1, Lcn2, Lcn4, Lcn5, Lcn6, Lcn7, and Lcn8 (Lcn3 was skipped). Later, these were named LC neurons, only unambiguous identities were maintained, and new numbers were given by (Otsuna and Ito, 2006). In Otsuna and Ito’s work, only Lcn4 and Lcn6 could be identified and became LC4 and LC6. However Lcn1, Lcn2, Lcn3, Lcn5, Lcn7, Lcn8 have no LC counterpart. In addition to LC4 and LC6, Otsuna and Ito identified LC9, LC10, LC11, LC12, LC13 and LC14. Naming of non-described types was based on the style of Otsuna and Ito (2006) and done in coordination with A. Nern and G. Rubin. Neuropils are referred to using the terminology of the Insect Brain Name Working Group (Ito et al., 2014). Abbreviations used: LC - lobula columnar; LPC - lobula plate columnar; LPLC - lobula plate, lobula columnar; MC - medulla columnar; Lat – lamina tangential. We call the union of the posterior ventrolateral protocerebrum (PVLP), posterior lateral protocerebrum (PLP) and anterior optic tubercle (AOTU) the optic Ventrolateral Neuropil (oVLNP).

## Acknowledgements

We thank Barry Dickson for access to the Vienna Tiles library and comments on the manuscript. We discussed with Aljosha Nern and Gerry Rubin a common nomenclature for the VPNs. We thank the Janelia Fly Light team and the Dickson lab for providing the datasets. IMP/IMBA Biooptics core facility provided extensive microscopy support. Flies were purchased from the Drosophila Bloomington Stock Center and the Vienna Drosophila RNAi Center. Arnim Jennet provided a 3D atlas of brain regions. Veit Grabe and Silke Sachse provided a 3D atlas of the antennal lobes. We thank Gaby Maimon and David Hain for comments on the manuscript. This work was supported by ERC Starting Grant 281884 “FlyVisualCircuits” to ADS, FFG Headquarter Grant 834223 to the IMP and VRVis, and by IMP core funding.

## Supplement Captions

**Figure 1-figure supplement 1.**
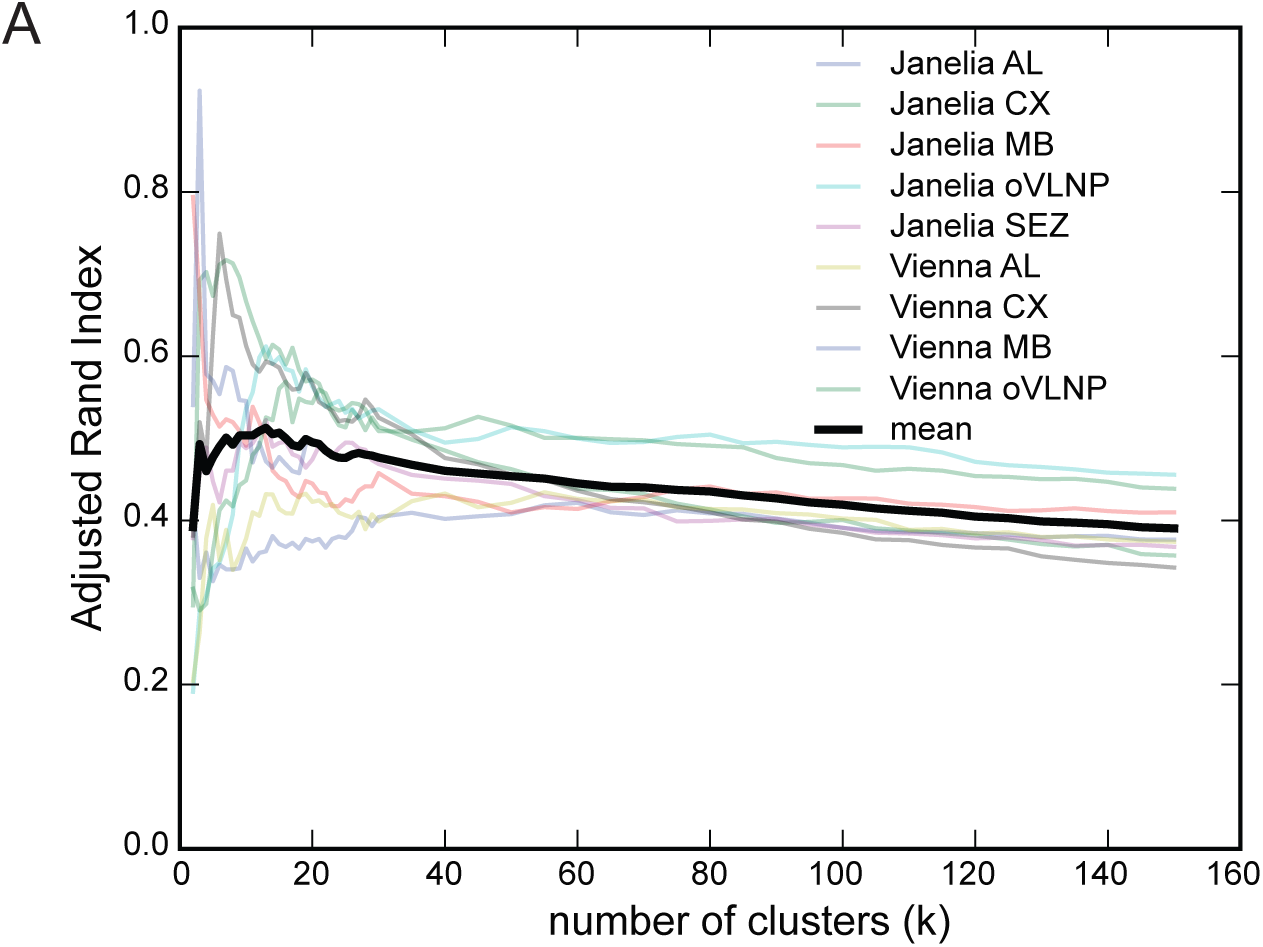
Repeatability scores across multiple runs of the *k*-medoids algorithm. The adjusted Rand index, a measure of repeatability, was calculated based on 10 repeated runs of the *k*-medoids algorithm for both datasets and several brain regions.

**Figure 2-figure supplement 1.**
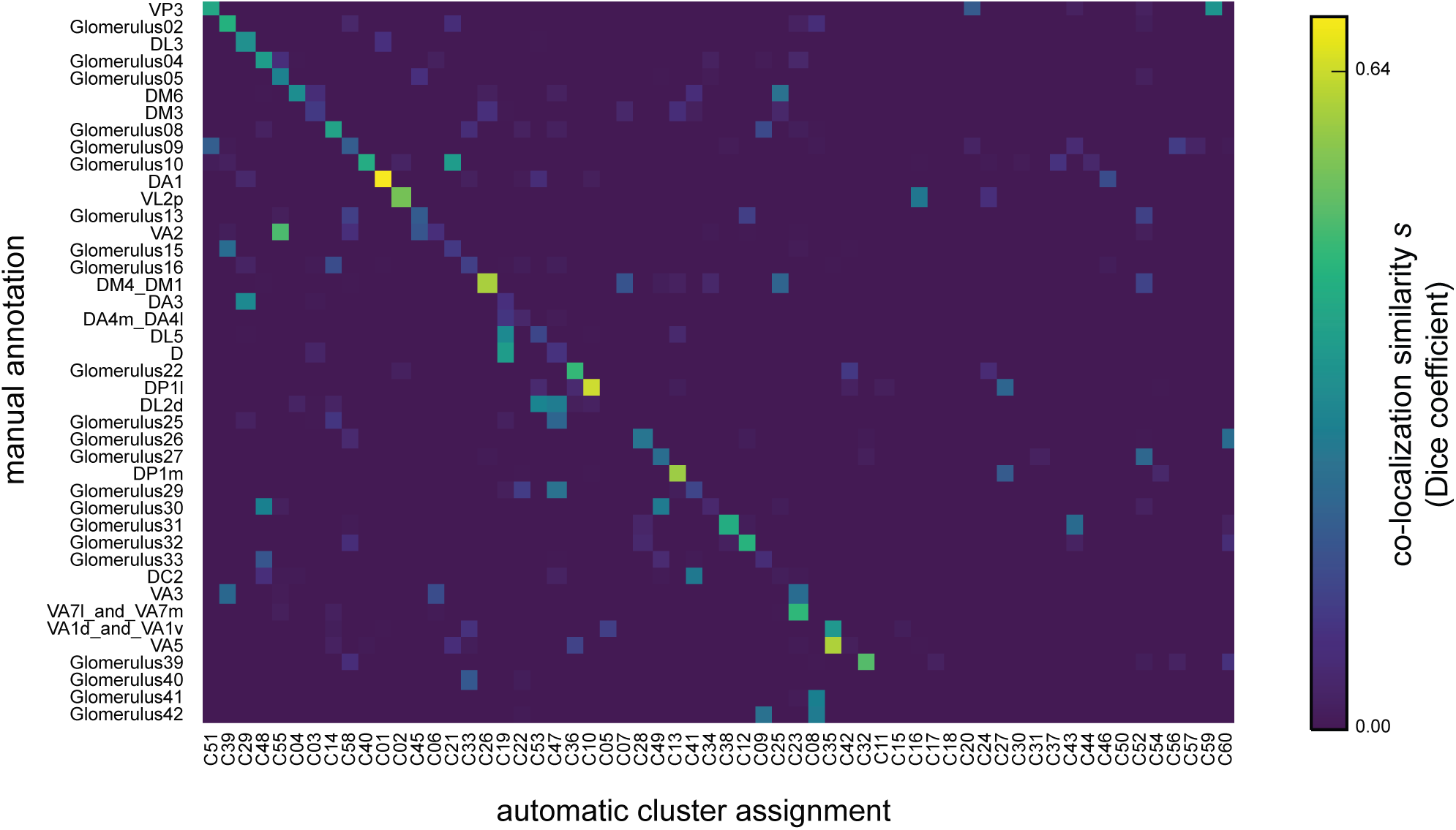
Automatically assigned clusters colocalize with manually segmented antennal lobe glomeruli. Colocalization similarity (measured based on set of voxels in manually annotated region and set of voxels in clustering result) between the Janelia FlyLight dataset and manual assignments using the same 3D template brain. (Janelia FlyLight data for the right antennal lobe region, run 1, 6502 voxels, 3462 driver lines, *k* equal 60.)

**Figure 2-figure supplement 2.**
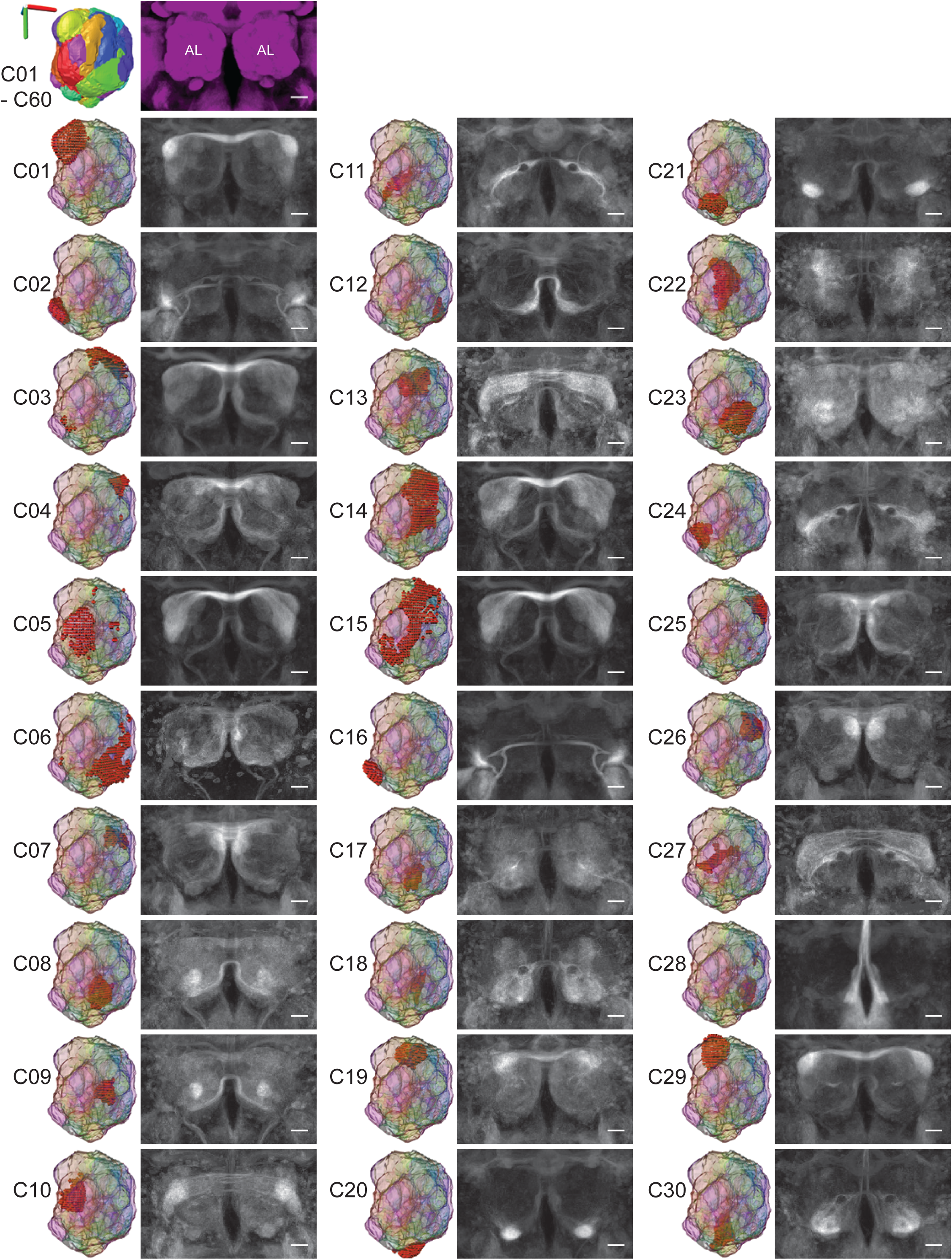
First 30 clusters from right antennal lobe. On the left of each column, a 3D rendering of each cluster is shown within the antennal lobe, and on the right is an average image of the drivers with high expression in that cluster but that do not broadly express. (Janelia FlyLight data for the right antennal lobe region, run 1, 6502 voxels, 3462 driver lines, *k* equal 60. Scale bars 20 μm.)

**Figure 2-figure supplement 3.**
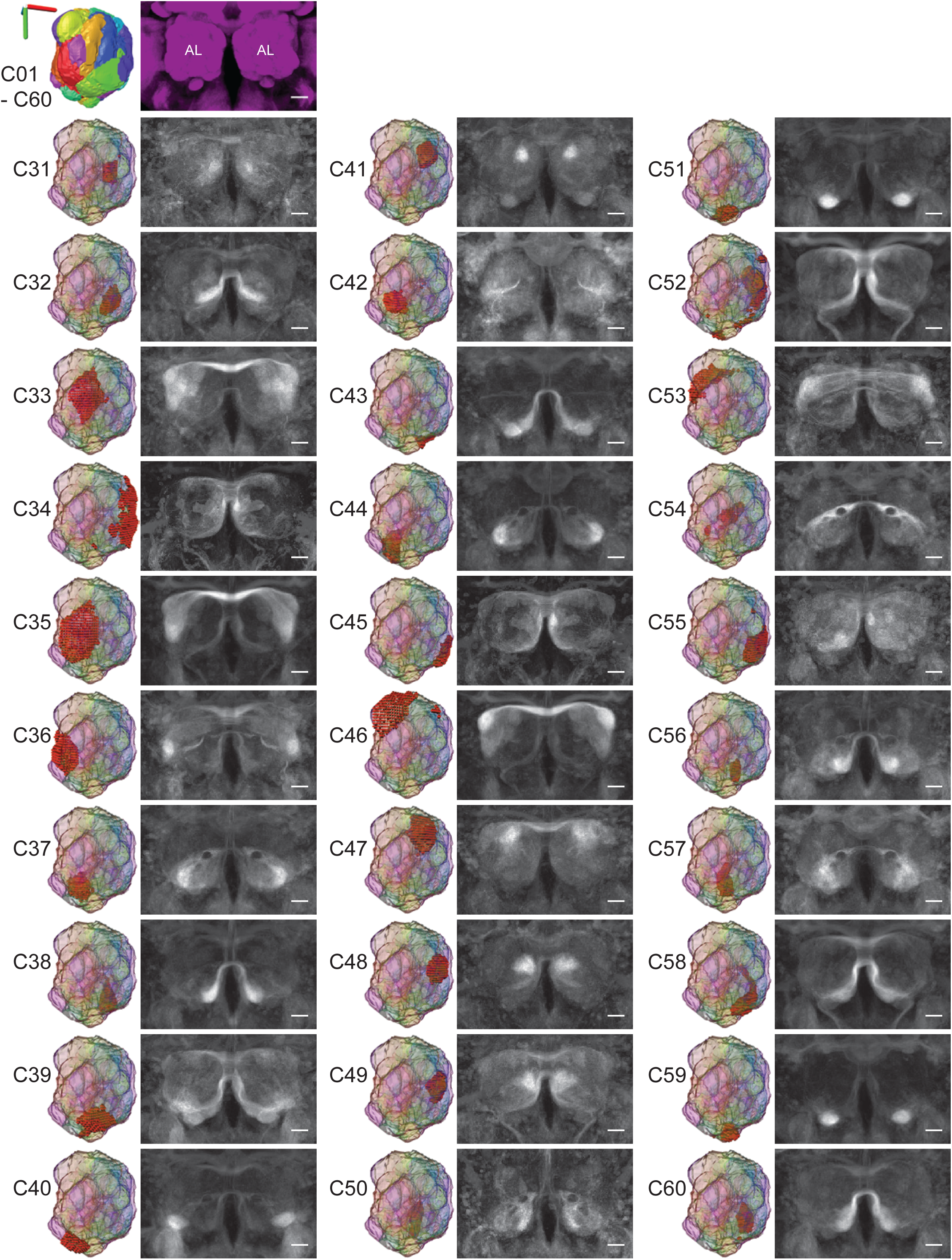
Second 30 clusters from right antennal lobe. As in Figure 2-figure supplement 2. (Janelia FlyLight data for the right antennal lobe region, run 1, 6502 voxels, 3462 driver lines, *k* equal 60. Scale bars 20 μm.

**Figure 2-figure supplement 4.**
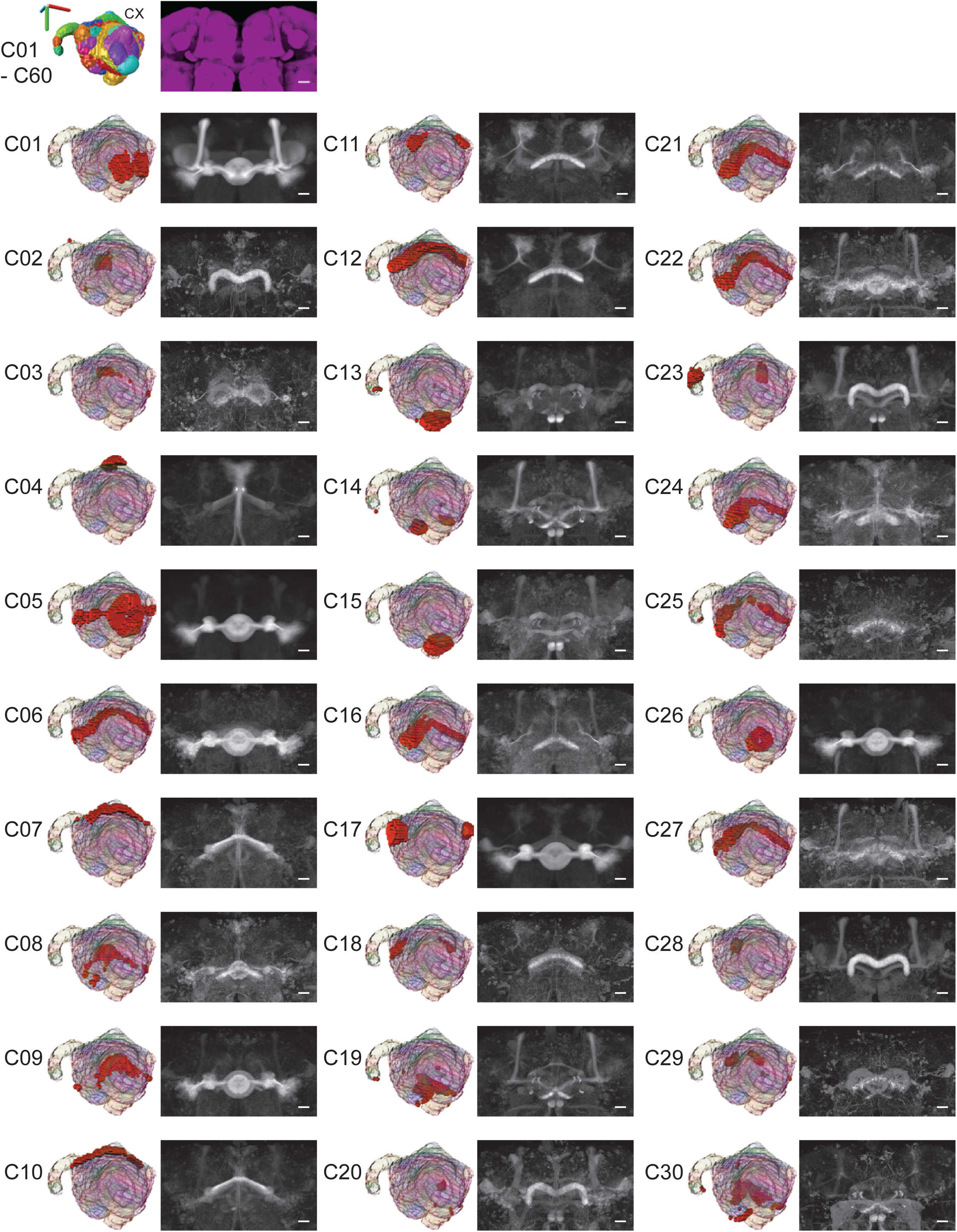
First 30 clusters from central complex. As in Figure 2-figure supplement 2 but for the central complex region. (Janelia FlyLight data for the central complex region, run 1, 27598 voxels, 3462 driver lines, *k* equal 60. Scale bars 20 μm.)

**Figure 2-figure supplement 5.**
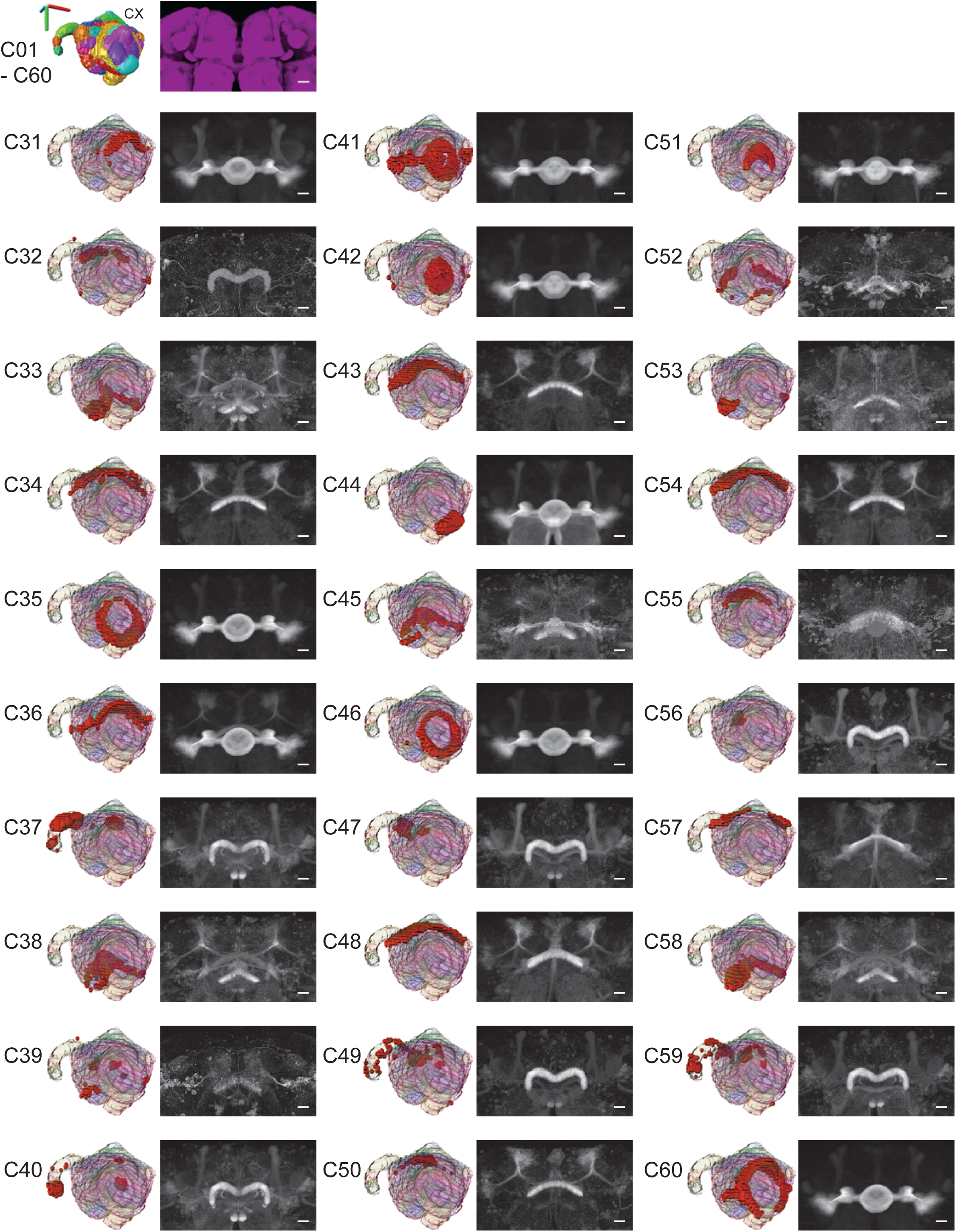
Second 30 clusters from central complex. As in Figure 2-figure supplement 4. (Janelia FlyLight data for the central complex region, run 1, 27598 voxels, 3462 driver lines, *k* equal 60. Scale bars 20 μm.)

**Figure 3-figure supplement 1.**
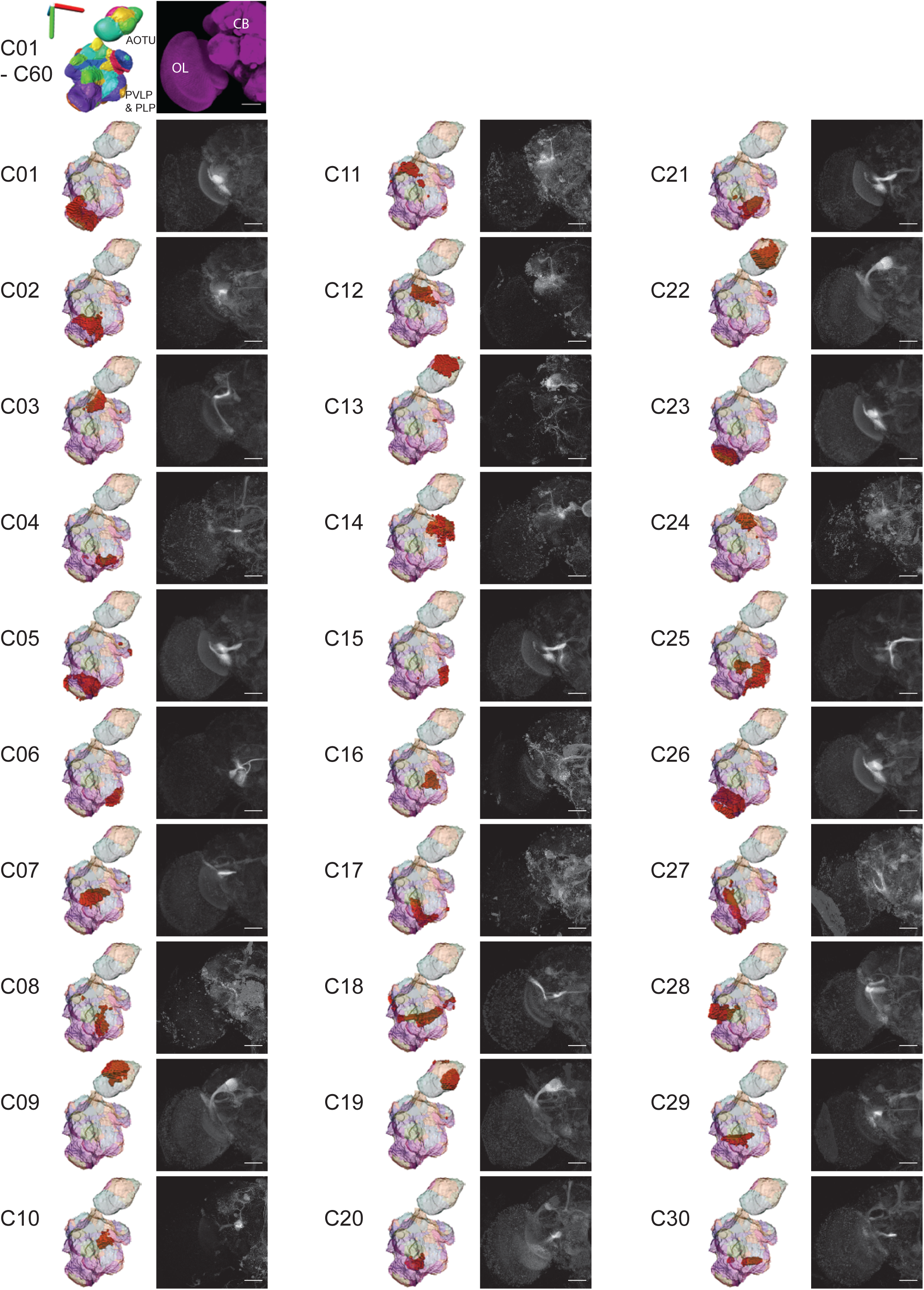
First 30 clusters from the oVLNP region, using Janelia FlyLight dataset. As in Figure 2-figure supplement 2 but for the oVLNP region. (Janelia FlyLight data for the oVLNP region defined as defined as PLP, PVLP, and AOTU, run 1, 42317 voxels, 3462 driver lines, *k* equal 60. Scale bars 50 μm.)

**Figure 3-figure supplement 2.**
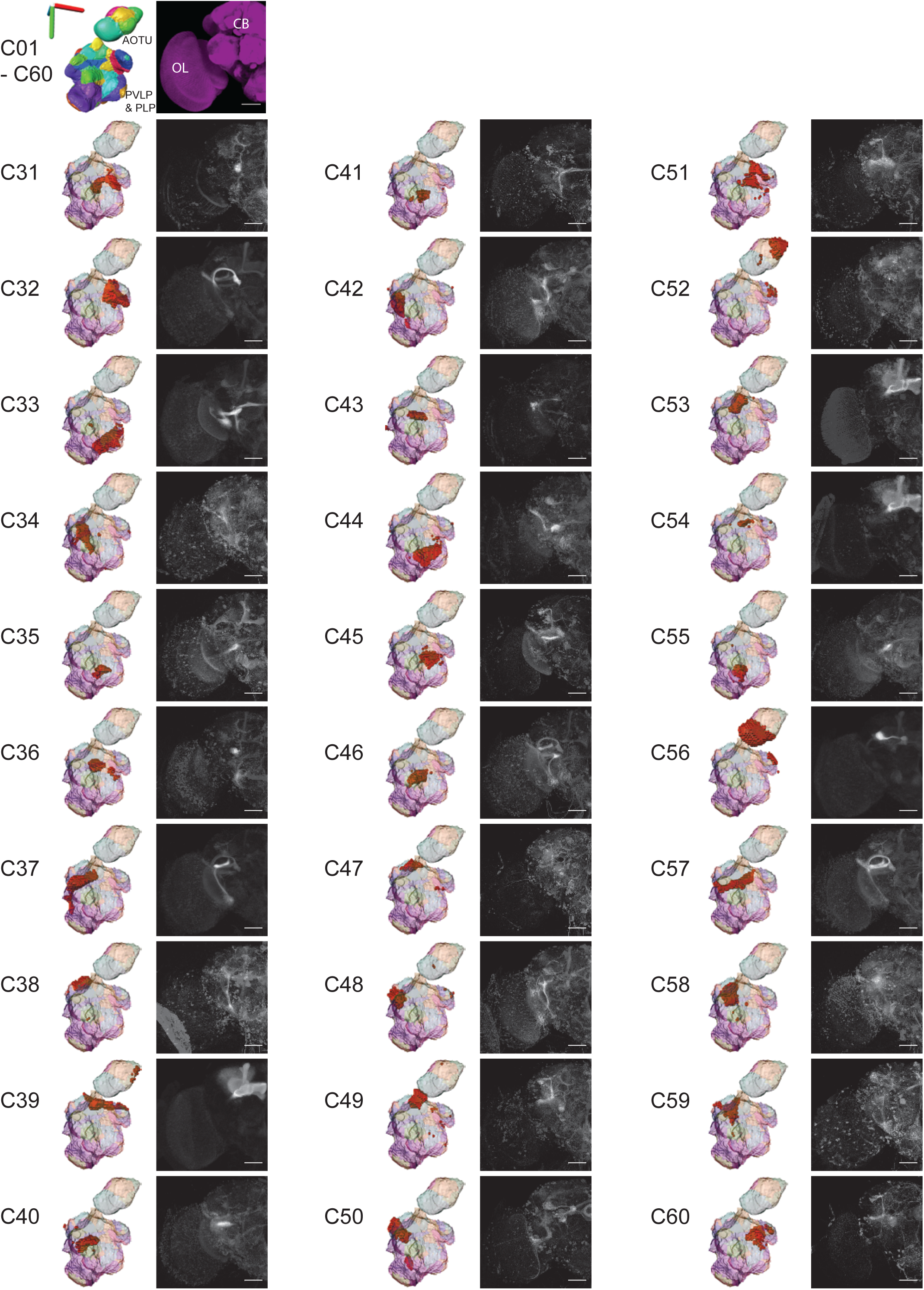
Second 30 clusters from the oVLNP region, using Janelia FlyLight dataset. As in Figure 3-figure supplement 1. (Janelia FlyLight data for the the oVLNP region defined as defined as PLP, PVLP, and AOTU, run 1, 42317 voxels, 3462 driver lines, *k* equal 60. Scale bars 50 μm.)

**Figure 4-figure supplement 1.**
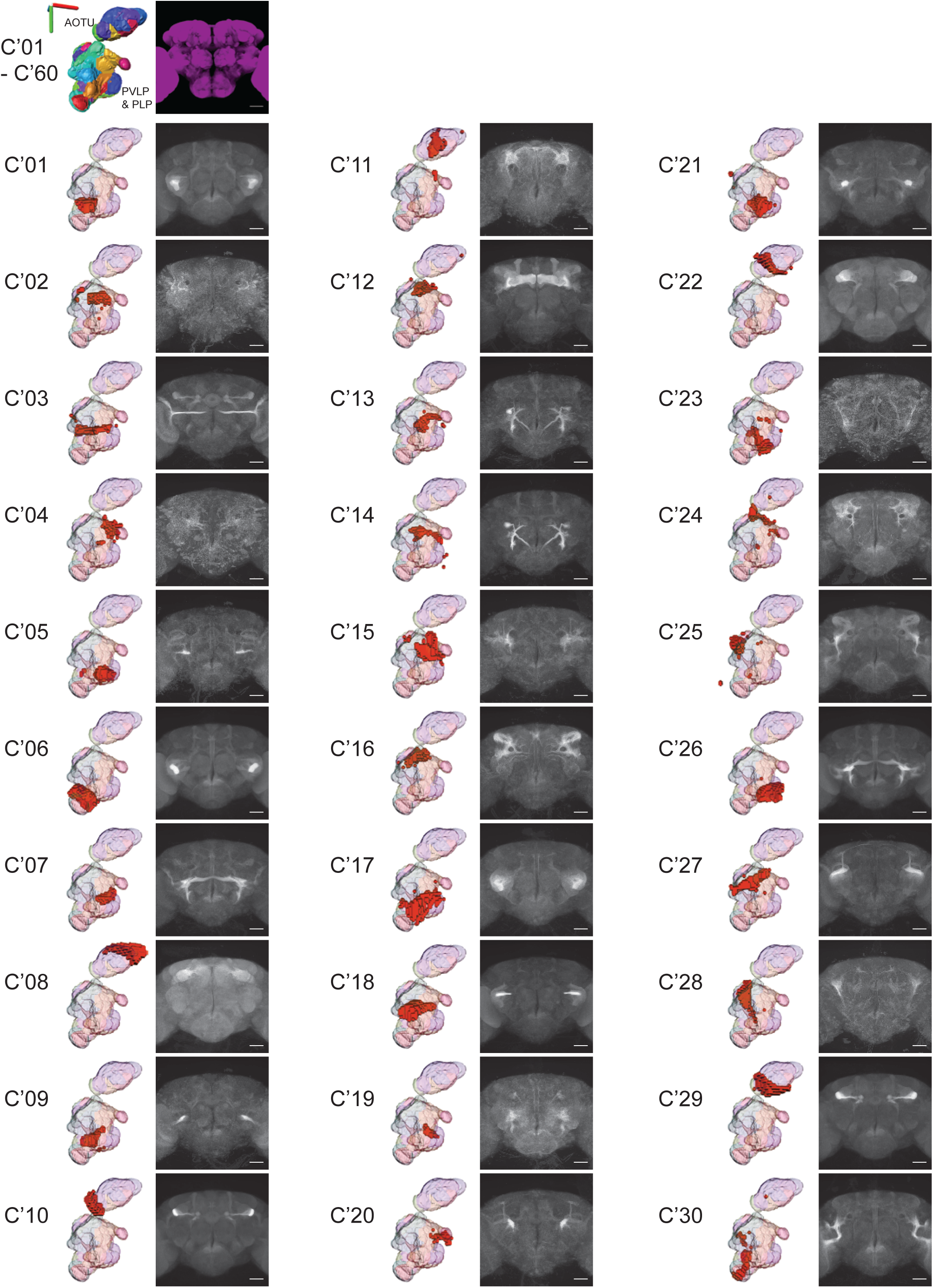
First 30 clusters from the oVLNP region, using Vienna Tiles dataset. As in Figure 3-figure supplement 1 but for the Vienna Tiles data. (Vienna Tiles data for the the oVLNP region defined as defined as PLP, PVLP, and AOTU, run 1, 13458 voxels, 6022 driver lines, *k* equal 60. Scale bars 50 μm.)

**Figure 4-figure supplement 2.**
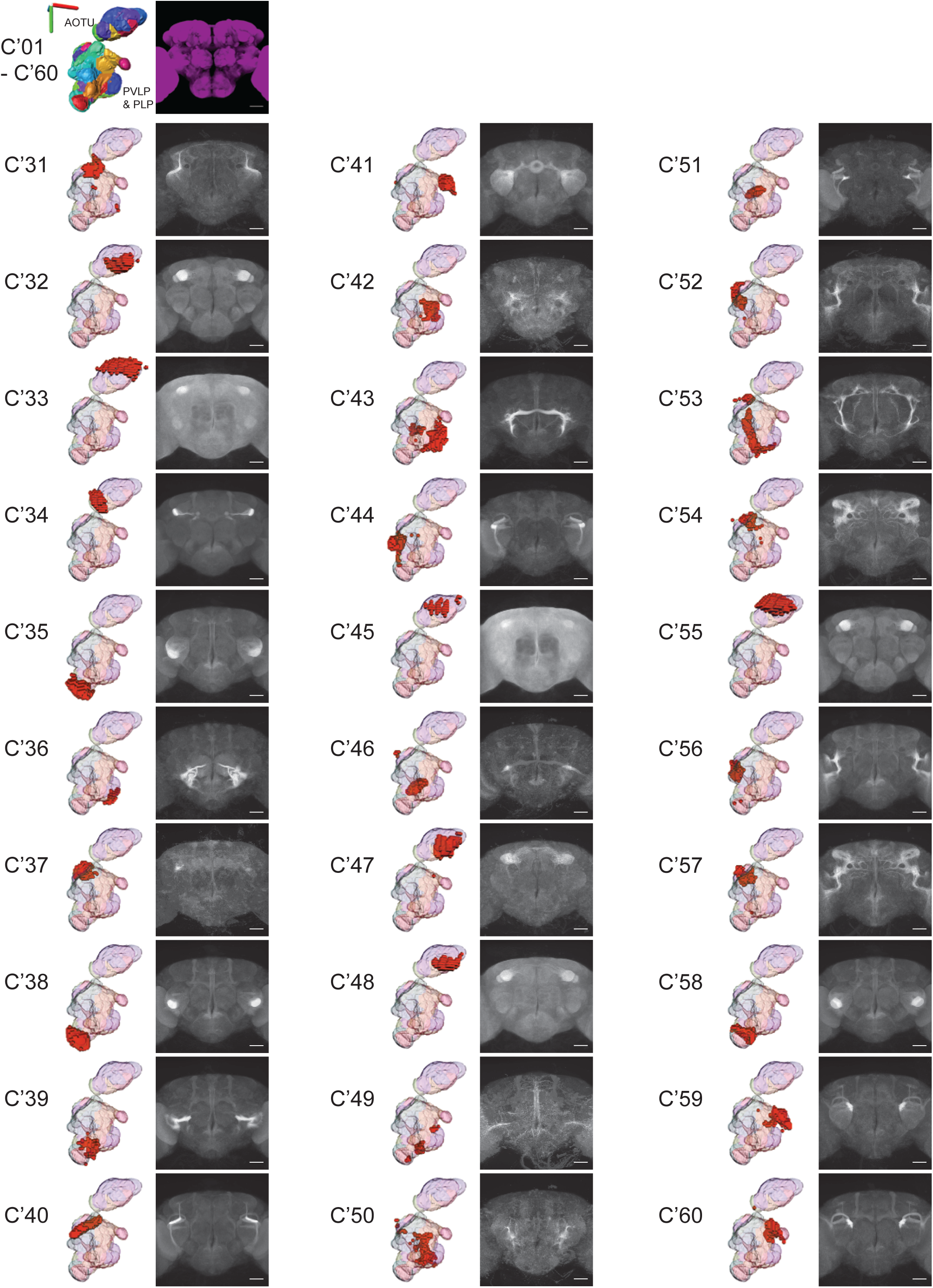
Second 30 clusters from the oVLNP region, using Vienna Tiles dataset. As in Figure 4-figure supplement 1. (Vienna Tiles data for the the oVLNP region defined as defined as PLP, PVLP, and AOTU, run 1, 13458 voxels, 6022 driver lines, *k* equal 60. Scale bars 50 μm.)

**Figure 7-figure supplement 1.**
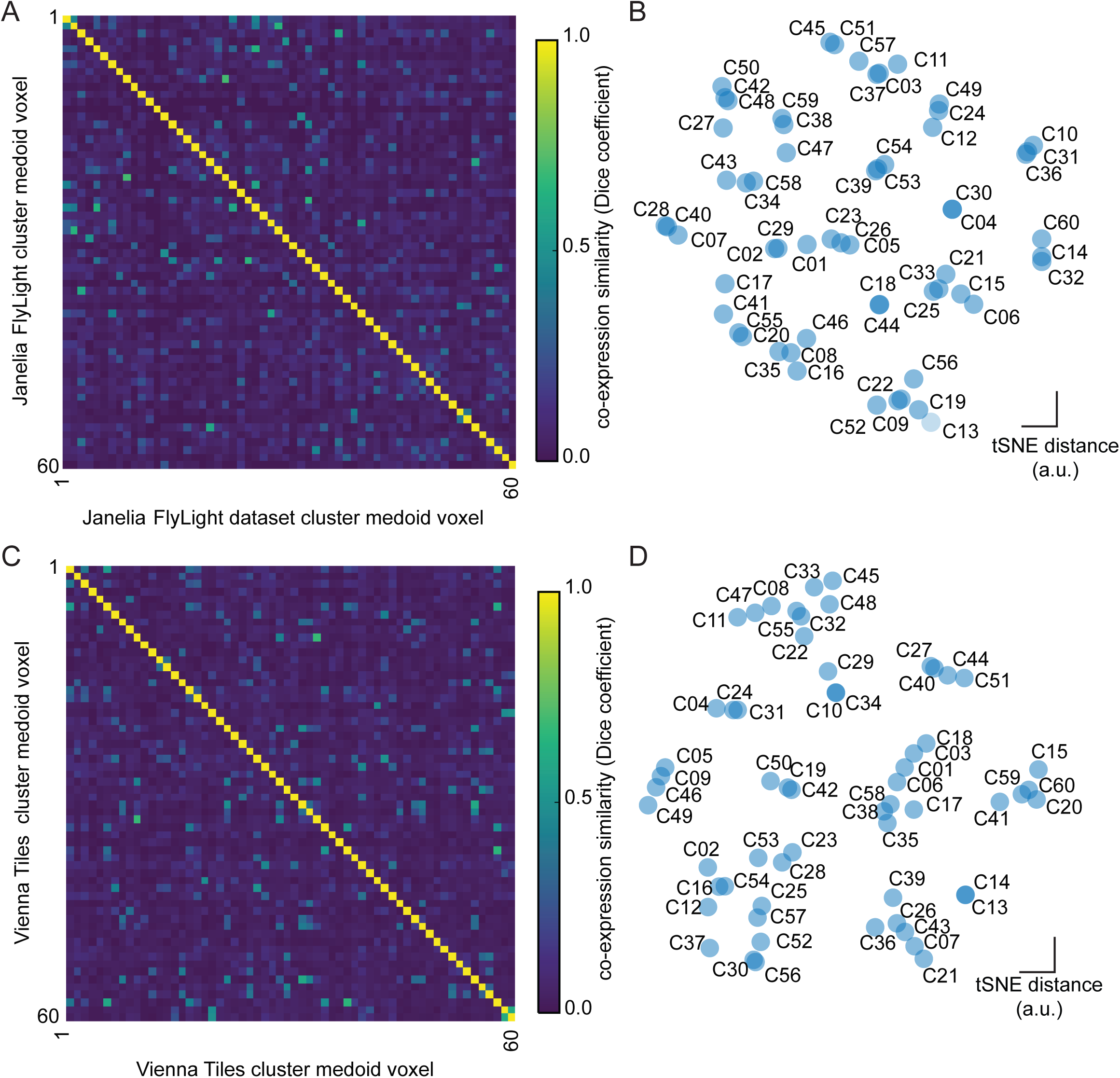
Clustering quality for both datasets. A) Quantification of similarity between clusters as measured by voxel-to-voxel similarity *s* for each medoid of every cluster of run 1 in the oVLNP region. B) t-distributed stochastic neighbor (tSNE) maps showing a representation of molecular distance between medoids in the oVLNP region of the Janelia FlyLight dataset. C) Quantification of similarity between clusters as measured by voxel-to-voxel similarity *s* for each medoid of every cluster in the oVLNP region of run 1 the Vienna Tiles dataset. D) t-distributed stochastic neighbor (tSNE) maps showing a representation of molecular distance between medoids in the oVLNP region of the Vienna Tiles dataset.

**Figure 7-figure supplement 2.**
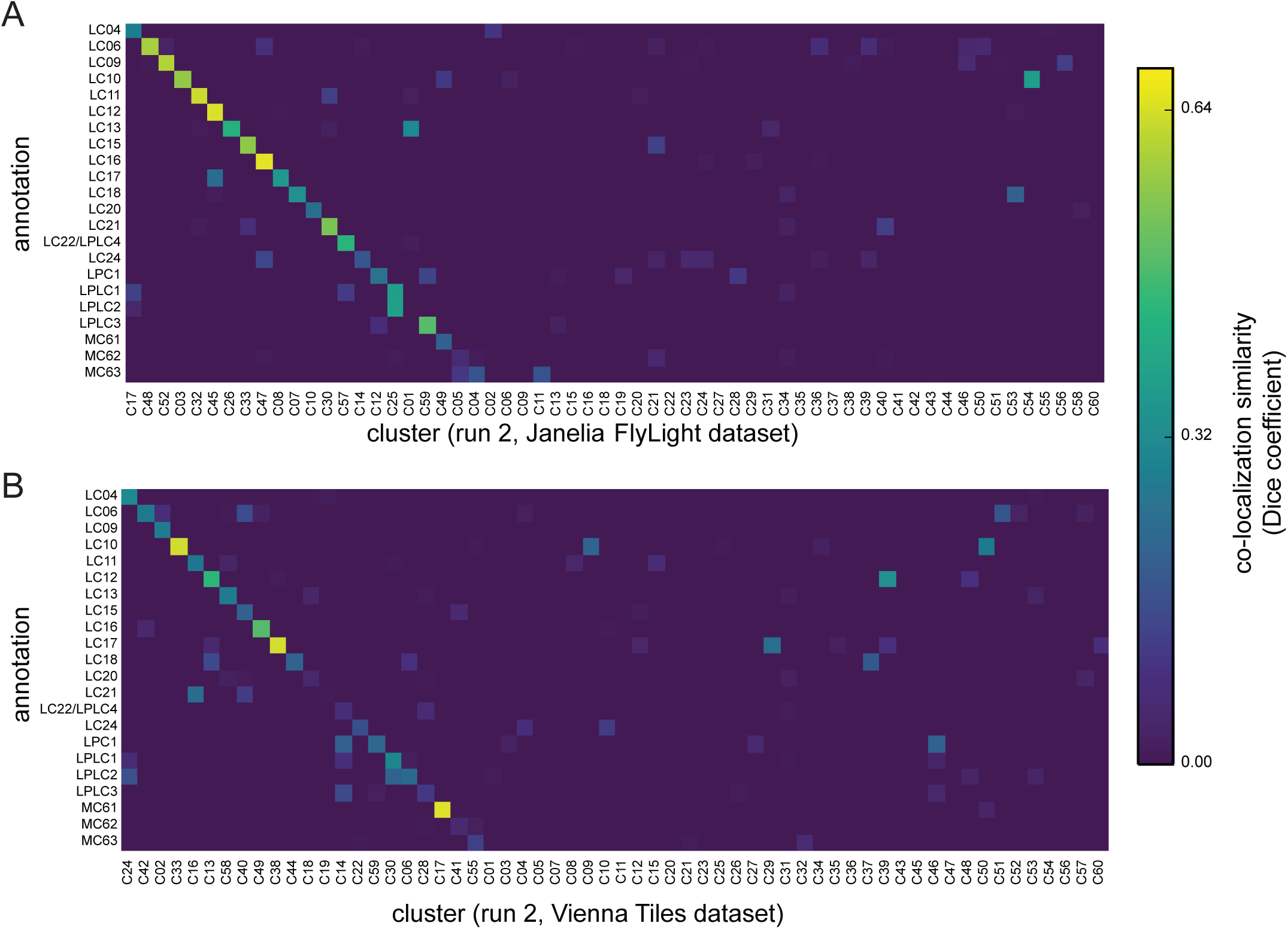
Repeated clustering of the same dataset gives similar results. A) Colocalization similarity (measured based on set of voxels in manually annotated region and set of voxels in clustering result) between a second clustering run on the Janelia FlyLight dataset and manual assignments using the same 3D template brain. Compare with Figure 7a. (Janelia FlyLight data for run 2, oVLNP, 42317 voxels, 3462 driver lines, *k* equal 60. Scale bars 50 μm.) B) Colocalization similarity between a second clustering run on the Vienna Tiles dataset and manual assignments using the same 3D template brain. (Vienna Tiles data for run 2, oVLNP, 13458 voxels, 6022 driver lines, *k* equal 60.)

**Figure 8-table supplement 1.**
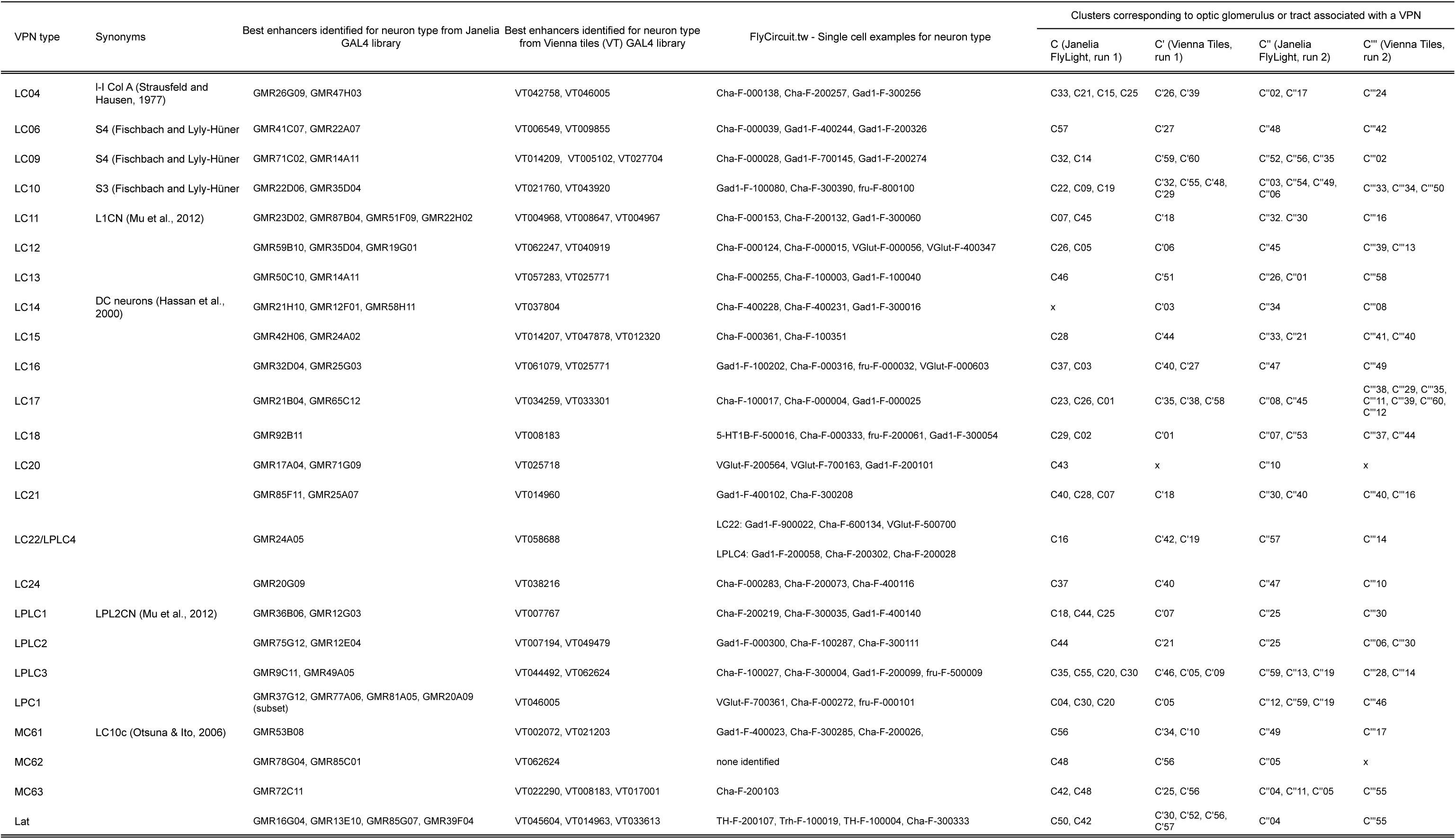
Table with VPN, Clusters, Driver lines, Flycircuit IDs.

## References

Ahrens, M.B., Li, J.M., Orger, M.B., Robson, D.N., Schier, A.F., Engert, F., Portugues, R., 2012. Brain-wide neuronal dynamics during motor adaptation in zebrafish. Nature. doi:10.1038/nature11057

Alkemade, A., Keuken, M.C., Forstmann, B.U., 2013. A perspective on terra incognita: uncovering the neuroanatomy of the human subcortex. Front. Neuroanat. 7. doi:10.3389/fnana.2013.00040

Alonso-Barba, J.I., Rahman, R.-U., Wittbrodt, J., Mateo, J.L., 2015. MEPD: medaka expression pattern database, genes and more. Nucleic Acids Res. gkv1029. doi:10.1093/nar/gkv1029

Aptekar, J.W., Keleş, M.F., Lu, P.M., Zolotova, N.M., Frye, M.A., 2015. Neurons forming optic glomeruli compute figure-ground discriminations in Drosophila. J. Neurosci. Off. J. Soc. Neurosci. 35, 7587–7599. doi:10.1523/JNEUROSCI.0652-15.2015

Bausenwein, B., Wolf, R., Heisenberg, M., 1986. Genetic Dissection of Optomotor Behavior in Drosophila melanogaster Studies on Wild-Type and the Mutant optomotor-blindH31. J. Neurogenet. 3, 87–109.

Bruckner, S., Soltészová, V., Gröller, M.E., Hladůvka, J., Bühler, K., Yu, J.Y., Dickson, B.J., 2009. BrainGazer-visual queries for neurobiology research. IEEE Trans. Vis. Comput. Graph. 15, 1497–504. doi:10.1109/TVCG.2009.121

Burkhardt, D., Motte, I.D., 1983. How Stalk-Eyed Flies Eye Stalk-Eyed Flies: Observations and Measurements of the Eyes of Cyrtodiopsis whitei (Diopsidae, Diptera). J Comp Physiol A 151, 407–421.

Cachero, S., Ostrovsky, A.D., Yu, J.Y., Dickson, B.J., Jefferis, G.S.X.E., 2010. Sexual Dimorphism in the Fly Brain. Curr. Biol. CB. doi:10.1016/j.cub.2010.07.045

Cardona, A., Saalfeld, S., Preibisch, S., Schmid, B., Cheng, A., Pulokas, J., Tomancak, P., Hartenstein, V., 2010. An integrated micro- and macroarchitectural analysis of the Drosophila brain by computer-assisted serial section electron microscopy. PLoS Biol. 8, 17. doi:10.1371/journal.pbio.1000502

Chiang, A.-S., Lin, C.-Y., Chuang, C.-C., Chang, H.-M., Hsieh, C.-H., Yeh, C.-W., Shih, C.-T., Wu, J.-J., Wang, G.-T., Chen, Y.-C., Wu, C.-C., Chen, G.-Y., Ching, Y.-T., Lee, P.-C., Lin, C.-Y., Lin, H.-H., Wu, C.-C., Hsu, H.-W., Huang, Y.-A., Chen, J.-Y., Chiang, H.-J., Lu, C.-F., Ni, R.-F., Yeh, C.-Y., Hwang, J.-K., 2011. Three-Dimensional Reconstruction of Brain-wide Wiring Networks in Drosophila at Single-Cell Resolution. Curr. Biol. 21, 1–11. doi:10.1016/j.cub.2010.11.056

Costa, M., Manton, J.D., Ostrovsky, A.D., Prohaska, S., Jefferis, G.S.X.E., 2015. NBLAST: Rapid, sensitive comparison of neuronal structure and construction of neuron family databases. bioRxiv 006346. doi:10.1101/006346

Couto, A., Alenius, M., Dickson, B.J., 2005. Molecular, Anatomical, and Functional Organization of the Drosophila Olfactory System. Curr. Biol. 15, 1535–1547. doi:10.1016/j.cub.2005.07.034

Fakhry, A., Ji, S., 2015. High-resolution prediction of mouse brain connectivity using gene expression patterns. Methods, Spatial mapping of multi-modal data in neuroscience 73, 71–78. doi:10.1016/j.ymeth.2014.07.011

Fischbach, K.-F., Dittrich, A., 1989. The optic lobe of Drosophila melanogaster. I. A Golgi analysis of wild-type structure. Cell Tissue Res. 258. doi:10.1007/BF00218858

Fischbach, K.-F., Lyly-Hünerberg, I., 1983. Genetic dissection of the anterior optic tract of Drosophila melanogaster. Cell Tissue Res 231, 551–563.

Goel, P., Kuceyeski, A., Locastro, E., Raj, A., 2014. Spatial patterns of genome-wide expression profiles reflect anatomic and fiber connectivity architecture of healthy human brain. Hum. Brain Mapp. 35, 4204–4218. doi:10.1002/hbm.22471

Grabe, V., Strutz, A., Baschwitz, A., Hansson, B.S., Sachse, S., 2015. Digital in vivo 3D atlas of the antennal lobe of Drosophila melanogaster. J. Comp. Neurol. 523, 530–544. doi:10.1002/cne.23697

Hadjieconomou, D., Rotkopf, S., Alexandre, C., Bell, D.M., Dickson, B.J., Salecker, I., 2011. Flybow: genetic multicolor cell labeling for neural circuit analysis in Drosophila melanogaster. Nat. Methods. doi:10.1038/nmeth.1567

Hampel, S., Chung, P., McKellar, C.E., Hall, D., Looger, L.L., Simpson, J.H., 2011. Drosophila Brainbow: a recombinase-based fluorescence labeling technique to subdivide neural expression patterns. Nat. Methods 8, 253–9. doi:10.1038/nmeth.1566

Hanesch, U., Fischbach, K.-F., Heisenberg, M., 1989. Neuronal architecture of the central complex in Drosophila melanogaster. Cell Tissue Res. 257, 343–366. doi:10.1007/BF00261838

Hawrylycz, M.J., Lein, E.S., Guillozet-Bongaarts, A.L., Shen, E.H., Ng, L., Miller, J.A., van de Lagemaat, L.N., Smith, K.A., Ebbert, A., Riley, Z.L., Abajian, C., Beckmann, C.F., Bernard, A., Bertagnolli, D., Boe, A.F., Cartagena, P.M., Chakravarty, M.M., Chapin, M., Chong, J., Dalley, R.A., Daly, B.D., Dang, C., Datta, S., Dee, N., Dolbeare, T.A., Faber, V., Feng, D., Fowler, D.R., Goldy, J., Gregor, B.W., Haradon, Z., Haynor, D.R., Hohmann, J.G., Horvath, S., Howard, R.E., Jeromin, A., Jochim, J.M., Kinnunen, M., Lau, C., Lazarz, E.T., Lee, C., Lemon, T.A., Li, L., Li, Y., Morris, J.A., Overly, C.C., Parker, P.D., Parry, S.E., Reding, M., Royall, J.J., Schulkin, J., Sequeira, P.A., Slaughterbeck, C.R., Smith, S.C., Sodt, A.J., Sunkin, S.M., Swanson, B.E., Vawter, M.P., Williams, D., Wohnoutka, P., Zielke, H.R., Geschwind, D.H., Hof, P.R., Smith, S.M., Koch, C., Grant, S.G.N., Jones, A.R., 2012. An anatomically comprehensive atlas of the adult human brain transcriptome. Nature 489, 391–399. doi:10.1038/nature11405

Helmstaedter, M., Briggman, K.L., Turaga, S.C., Jain, V., Seung, H.S., Denk, W., 2013. Connectomic reconstruction of the inner plexiform layer in the mouse retina. Nature 500, 168–174. doi:10.1038/nature12346

Ito, K., Shinomiya, K., Ito, M., Armstrong, J.D., Boyan, G., Hartenstein, V., Harzsch, S., Heisenberg, M., Homberg, U., Jenett, A., Keshishian, H., Restifo, L.L., Rössler, W., Simpson, J.H., Strausfeld, N.J., Strauss, R., Vosshall, L.B., 2014. A Systematic Nomenclature for the Insect Brain. Neuron 81, 755–765. doi:10.1016/j.neuron.2013.12.017

Ito, M., Masuda, N., Shinomiya, K., Endo, K., Ito, K., 2013. Systematic Analysis of Neural Projections Reveals Clonal Composition of the Drosophila Brain. Curr. Biol. 1–12. doi:10.1016/j.cub.2013.03.015

Jenett, A., Rubin, G.M., Ngo, T.-T.B., Shepherd, D., Murphy, C., Dionne, H., Pfeiffer, B.D., Cavallaro, A., Hall, D., Jeter, J., Iyer, N., Fetter, D., Hausenfluck, J.H., Peng, H., Trautman, E.T., Svirskas, R.R., Myers, E.W., Iwinski, Z.R., Aso, Y., DePasquale, G.M., Enos, A., Hulamm, P., Lam, S.C.B., Li, H.-H., Laverty, T.R., Long, F., Qu, L., Murphy, S.D., Rokicki, K., Safford, T., Shaw, K., Simpson, J.H., Sowell, A., Tae, S., Yu, Y., Zugates, C.T., 2012. A GAL4-driver line resource for Drosophila neurobiology. Cell Rep. 2, 991–1001. doi:10.1016/j.celrep.2012.09.011

Kaufman, L., Rousseeuw, P.J., 1987. Clustering by Means of Medoids, in: Statistical Data Analysis Based on the L1 Norm and Related Methods.

Kawakami, K., Abe, G., Asada, T., Asakawa, K., Fukuda, R., Ito, A., Lal, P., Mouri, N., Muto, A., Suster, M.L., Takakubo, H., Urasaki, A., Wada, H., Yoshida, M., 2010. zTrap: zebrafish gene trap and enhancer trap database. BMC Dev. Biol. 10, 105. doi:10.1186/1471-213X-10-105

Kondrychyn, I., Teh, C., Garcia-Lecea, M., Guan, Y., Kang, A., Korzh, V., 2011. Zebrafish Enhancer TRAP transgenic line database ZETRAP 2.0. Zebrafish 8, 181–182. doi:10.1089/zeb.2011.0718

Kubo, F., Hablitzel, B., Dal Maschio, M., Driever, W., Baier, H., Arrenberg, A.B., 2014. Functional Architecture of an Optic Flow-Responsive Area that Drives Horizontal Eye Movements in Zebrafish. Neuron 81, 1344–1359. doi:10.1016/j.neuron.2014.02.043

Kvon, E.Z., Kazmar, T., Stampfel, G., Yáñnez-Cuna, J.O., Pagani, M., Schernhuber, K., Dickson, B.J., Stark, A., 2014. Genome-scale functional characterization of Drosophila developmental enhancers in vivo. Nature. doi:10.1038/nature13395

Lein, E.S., Hawrylycz, M.J., Ao, N., Ayres, M., Bensinger, A., Bernard, A., Boe, A.F., Boguski, M.S., Brockway, K.S., Byrnes, E.J., Chen, L., Chen, L., Chen, T.-M., Chi Chin, M., Chong, J., Crook, B.E., Czaplinska, A., Dang, C.N., Datta, S., Dee, N.R., Desaki, A.L., Desta, T., Diep, E., Dolbeare, T.A., Donelan, M.J., Dong, H.-W., Dougherty, J.G., Duncan, B.J., Ebbert, A.J., Eichele, G., Estin, L.K., Faber, C., Facer, B.A., Fields, R., Fischer, S.R., Fliss, T.P., Frensley, C., Gates, S.N., Glattfelder, K.J., Halverson, K.R., Hart, M.R., Hohmann, J.G., Howell, M.P., Jeung, D.P., Johnson, R.A., Karr, P.T., Kawal, R., Kidney, J.M., Knapik, R.H., Kuan, C.L., Lake, J.H., Laramee, A.R., Larsen, K.D., Lau, C., Lemon, T.A., Liang, A.J., Liu, Y., Luong, L.T., Michaels, J., Morgan, J.J., Morgan, R.J., Mortrud, M.T., Mosqueda, N.F., Ng, L.L., Ng, R., Orta, G.J., Overly, C. C., Pak, T.H., Parry, S.E., Pathak, S.D., Pearson, O.C., Puchalski, R.B., Riley, Z.L., Rockett, H.R., Rowland, S.A., Royall, J.J., Ruiz, M.J., Sarno, N.R., Schaffnit, K., Shapovalova, N.V., Sivisay, T., Slaughterbeck, C.R., Smith, S.C., Smith, K.A., Smith, B.I., Sodt, A.J., Stewart, N.N., Stumpf, K.-R., Sunkin, S.M., Sutram, M., Tam, A., Teemer, C.D., Thaller, C., Thompson, C.L., Varnam, L.R., Visel, A., Whitlock, R.M., Wohnoutka, P.E., Wolkey, C.K., Wong, V.Y., Wood, M., Yaylaoglu, M.B., Young, R.C., Youngstrom, B.L., Feng Yuan, X., Zhang, B., Zwingman, T.A., Jones, A.R., 2007. Genome-wide atlas of gene expression in the adult mouse brain. Nature 445, 168–176. doi:10.1038/nature05453

Lin, C.-Y., Chuang, C.-C., Hua, T.-E., Chen, C.-C., Dickson, B.J., Greenspan, R.J., Chiang, A.-S., 2013. A comprehensive wiring diagram of the protocerebral bridge for visual information processing in the Drosophila brain. Cell Rep. 3, 1739–53. doi:10.1016/j.celrep.2013.04.022

Lister, J.A., 2011. Use of phage φC31 integrase as a tool for zebrafish genome manipulation. Methods Cell Biol. 104, 195–208. doi:10.1016/B978-0-12-374814-0.00011-2

Livet, J., Weissman, T.A., Kang, H., Draft, R.W., Lu, J., Bennis, R.A., Sanes, J.R., Lichtman, J.W., 2007. Transgenic strategies for combinatorial expression of fluorescent proteins in the nervous system. Nature 450, 56–62. doi:10.1038/nature06293

Mahfouz, A., van de Giessen, M., van der Maaten, L., Huisman, S., Reinders, M., Hawrylycz, M.J. Lelieveldt, B.P.F., 2015. Visualizing the spatial gene expression organization in the brain through non-linear similarity embeddings. Methods, Spatial mapping of multi-modal data in neuroscience 73, 79–89. doi:10.1016/j.ymeth.2014.10.004

Masse, N.Y., Cachero, S., Ostrovsky, A., Jefferis, G.S.X.E., 2012. A mutual information approach to automate identification of neuronal clusters in Drosophila brain images. Front. Neuroinformatics 6, 21. doi:10.3389/fninf.2012.00021

Mosimann, C., Puller, A.-C., Lawson, K.L., Tschopp, P., Amsterdam, A., Zon, L.I., 2013. Site-directed zebrafish transgenesis into single landing sites with the phiC31 integrase system. Dev. Dyn. Off. Publ. Am. Assoc. Anat. 242, 949–963. doi:10.1002/dvdy.23989

Mu, L., Ito, K., Bacon, J.P., Strausfeld, N.J., 2012. Optic glomeruli and their inputs in Drosophila share an organizational ground pattern with the antennal lobes. J. Neurosci. Off. J. Soc. Neurosci. 32, 6061–71. doi:10.1523/JNEUROSCI.0221-12.2012

Myers, E.M., Bartlett, C.W., Machiraju, R., Bohland, J.W., 2015. An integrative analysis of regional gene expression profiles in the human brain. Methods, Spatial mapping of multi-modal data in neuroscience 73, 54–70. doi:10.1016/j.ymeth.2014.12.010

Nern, A., Pfeiffer, B.D., Rubin, G.M., 2015. Optimized tools for multicolor stochastic labeling reveal diverse stereotyped cell arrangements in the fly visual system. Proc. Natl. Acad. Sci. U. S. A. doi:10.1073/pnas.1506763112

Ng, L., Bernard, A., Lau, C., Overly, C.C., Dong, H.-W., Kuan, C., Pathak, S., Sunkin, S.M., Dang, C., Bohland, J.W., Bokil, H., Mitra, P.P., Puelles, L., Hohmann, J., Anderson, D.J., Lein, E.S., Jones, A.R., Hawrylycz, M., 2009. An anatomic gene expression atlas of the adult mouse brain. Nat. Neurosci. 12, 356–362. doi:10.1038/nn.2281

Nicolaï, L.J.J., Ramaekers, A., Raemaekers, T., Drozdzecki, A., Mauss, A.S., Yan, J., Landgraf, M., Annaert, W., Hassan, B.A., 2010. Genetically encoded dendritic marker sheds light on neuronal connectivity in Drosophila. Proc. Natl. Acad. Sci. U. S. A. 107, 20553–20558. doi:10.1073/pnas.1010198107

Okamura, J.-Y., Strausfeld, N.J., 2007. Visual system of calliphorid flies: motion- and orientation-sensitive visual interneurons supplying dorsal optic glomeruli. J. Comp. Neurol. 500, 189–208. doi:10.1002/cne.21195

Otsuna, H., Ito, K., 2006. Systematic analysis of the visual projection neurons of Drosophila melanogaster. I. Lobula-specific pathways. J. Comp. Neurol. 497, 928–58. doi:10.1002/cne.21015

Otsuna, H., Shinomiya, K., Ito, K., 2014. Parallel neural pathways in higher visual centers of the Drosophila brain that mediate wavelength-specific behavior. Front. Neural Circuits 8, 8. doi:10.3389/fncir.2014.00008

Pedregosa, F., Varoquaux, G., Gramfort, A., Michel, V., Thirion, B., Grisel, O., Blondel, M., Prettenhofer, P., Weiss, R., Dubourg, V., Vanderplas, J., Passos, A., Cournapeau, D., Brucher, M., Perrot, M., Duchesnay, E., 2011. Scikit-learn: machine learning in Python. J Mach Learn Res.

Pfeiffer, B.D., Jenett, A., Hammonds, A.S., Ngo, T.-T.B., Misra, S., Murphy, C., Scully, A., Carlson, J.W., Wan, K.H., Laverty, T.R., Mungall, C., Svirskas, R., Kadonaga, J.T., Doe, C.Q., Eisen, M.B., Celniker, S.E., Rubin, G.M., 2008. Tools for neuroanatomy and neurogenetics in Drosophila. PNAS 105, 9715–20. doi:10.1073/pnas.0803697105

Pfeiffer, B.D., Ngo, T.-T.B., Hibbard, K.L., Murphy, C., Jenett, A., Truman, J.W., Rubin, G.M., 2010. Refinement of Tools for Targeted Gene Expression in Drosophila. Genetics 186, 735–755. doi:10.1534/genetics.110.119917

Phelan, P., Nakagawa, M., Wilkin, M.B., Moffat, K.G., O’Kane, C.J., Davies, J.A., Bacon, J.P., 1996. Mutations Drosophila in shaking-B Prevent Giant Fiber System Electrical Synapse Formation in the Drosophila Giant Fiber System. J Neurosci 16, 1101–1113.

Portugues, R., Feierstein, C.E., Engert, F., Orger, M.B., 2014. Article Whole-Brain Activity Maps Reveal Stereotyped, Distributed Networks for Visuomotor Behavior. Neuron 81, 1328–1343. doi:10.1016/j.neuron.2014.01.019

Raghu, S.V., Borst, A., 2011. Candidate Glutamatergic Neurons in the Visual System of Drosophila. PLoS ONE 6, e19472. doi:10.1371/journal.pone.0019472

Raghu, S.V., Joesch, M., Borst, A., Reiff, D.F., 2007. Synaptic organization of lobula plate tangential cells in Drosophila: gamma-aminobutyric acid receptors and chemical release sites. J. Comp. Neurol. 502, 598–610. doi:10.1002/cne.21319

Raghu, S.V., Joesch, M., Sigrist, S.J., Borst, A., Reiff, D.F., 2009. Synaptic organization of lobula plate tangential cells in Drosophila: Dalpha7 cholinergic receptors. J. Neurogenet. 23, 200–9. doi:10.1080/01677060802471684

Raghu, S.V., Reiff, D.F., Borst, A., 2011. Neurons with cholinergic phenotype in the visual system of Drosophila. J. Comp. Neurol. 519, 162–76. doi:10.1002/cne.22512

Randlett, O., Wee, C.L., Naumann, E.A., Nnaemeka, O., Schoppik, D., Fitzgerald, J.E., Portugues, R., Lacoste, A.M.B., Riegler, C., Engert, F., Schier, A.F., 2015. Whole-brain activity mapping onto a zebrafish brain atlas. Nat. Methods advance online publication. doi:10.1038/nmeth.3581

Ronneberger, O., Liu, K., Rath, M., Rueβ, D., Mueller, T., Skibbe, H., Drayer, B., Schmidt, T., Filippi, A., Nitschke, R., Brox, T., Burkhardt, H., Driever, W., 2012. ViBE-Z: a framework for 3D virtual colocalization analysis in zebrafish larval brains. Nat. Methods 9, 735–742. doi:10.1038/nmeth.2076

Shih, C.-T., Sporns, O., Yuan, S.-L., Su, T.-S., Lin, Y.-J., Chuang, C.-C., Wang, T.-Y., Lo, C.-C., Greenspan, R.J., Chiang, A.-S., 2015. Connectomics-Based Analysis of Information Flow in the Drosophila Brain. Curr. Biol. 25, 1249–1258. doi:10.1016/j.cub.2015.03.021

Strausfeld, N.J., Bacon, J.P., 1983. Multimodal convergence in the central nervous system of dipterous insects, in: Fortschritteder Zoologie: Multimodal Convergence in Sensory Systems. Gustav Fischer Verlag, New York, pp. 47–76.

Strausfeld, N.J., Lee, J.-K., 1991. Neuronal basis for parallel visual processing in the fly. Vis. Neurosci. 7, 13–33.

Strausfeld, N.J., Okamura, J.-Y., 2007. Visual system of calliphorid flies: organization of optic glomeruli and their lobula complex efferents. J. Comp. Neurol. 500, 166–88. doi:10.1002/cne.21196

Strausfeld, N.J., Sinakevitch, I., Okamura, J.-Y., 2007. Organization of local interneurons in optic glomeruli of the dipterous visual system and comparisons with the antennal lobes. Dev. Neurobiol. 67, 1267–88. doi:10.1002/dneu.20396

Strauss, R., Heisenberg, M., 1993. A higher control center of locomotor behavior in the Drosophila brain. J Neurosci 13, 1852–1861.

Takemura, S., Bharioke, A., Lu, Z., Nern, A., Vitaladevuni, S., Rivlin, P.K., Katz, W.T., Olbris, D.J., Plaza, S.M., Winston, P., Zhao, T., Horne, J.A., Fetter, R.D., Takemura, S., Blazek, K., Chang, L.-A., Ogundeyi, O., Saunders, M. a., Shapiro, V., Sigmund, C., Rubin, G.M., Scheffer, L.K., Meinertzhagen, I. a., Chklovskii, D.B., 2013. A visual motion detection circuit suggested by Drosophila connectomics. Nature 500, 175–181. doi:10.1038/nature12450

Thompson, C.L., Ng, L., Menon, V., Martinez, S., Lee, C.-K., Glattfelder, K., Sunkin, S.M., Henry, A., Lau, C., Dang, C., Garcia-Lopez, R., Martinez-Ferre, A., Pombero, A., Rubenstein, J.L.R., Wakeman, W.B., Hohmann, J., Dee, N., Sodt, A.J., Young, R., Smith, K., Nguyen, T.-N., Kidney, J., Kuan, L., Jeromin, A., Kaykas, A., Miller, J., Page, D., Orta, G., Bernard, A., Riley, Z., Smith, S., Wohnoutka, P., Hawrylycz, M.J., Puelles, L., Jones, A.R., 2014. A High-Resolution Spatiotemporal Atlas of Gene Expression of the Developing Mouse Brain. Neuron 83, 309–323. doi:10.1016/j.neuron.2014.05.033

von der Maaten, L.J.P., Hinton, G.E., 2008. Visualizing Data using t-SNE. J. Mach. Learn. Res.

Vosshall, L.B., Wong, A.M., Axel, R., 2000. An Olfactory Sensory Map in the Fly Brain. Cell 102, 147–159. doi:10.1016/S0092-8674(00)00021-0

White, J.G., Southgate, E., Thomson, J.N., Brenner, S., 1986. The Structure of the Nervous System of the Nematode Caenorhabditis elegans. Philos. Trans. R. Soc. B Biol. Sci. 314, 1–340. doi:10.1098/rstb.1986.0056

Wolff, T., Iyer, N.A., Rubin, G.M., 2015. Neuroarchitecture and neuroanatomy of the Drosophila central complex: A GAL4-based dissection of protocerebral bridge neurons and circuits. J. Comp. Neurol. 523, Spc1–Spc1. doi:10.1002/cne.23773

Yu, H., Awasaki, T., Schroeder, M.D.D., Long, F., Yang, J.S.S., He, Y., Ding, P., Kao, J., Wu, G.Y.-Y.Y., Peng, H., Myers, G., Lee, T., 2013. Clonal Development and Organization of the Adult Drosophila Central Brain. Curr Biol 23, 1–11. doi:10.1016/j.cub.2013.02.057

Yu, J.Y., Kanai, M.I., Demir, E., Jefferis, G.S.X.E., Dickson, B.J., 2010. Cellular Organization of the Neural Circuit that Drives Drosophila Courtship Behavior. Curr Biol 20, 1602–1614. doi:10.1016/j.cub.2010.08.025

Zhang, Y.Q., Rodesch, C.K., Broadie, K., 2002. Living synaptic vesicle marker: synaptotagmin-GFP. Genes. N. Y. N 2000 34, 142–145. doi:10.1002/gene.10144

